# Tonsillar expression quantitative trait loci verify and expand genetic contributors to childhood atopic diseases

**DOI:** 10.64898/2026.03.13.711667

**Authors:** Kim Lorenz, Samuel Yoon, Carole Le Coz, Karen Zur, Andrew Wells, Neil Romberg, Benjamin F. Voight

## Abstract

The spectrum of causal variants, mechanisms, and immunologic gene networks that influence pediatric atopic traits are not completely understood. Human genetic variation associated with transcript abundance (eQTLs) can help to advance our understanding, yet prior work has focused on profiling immune cell populations collected from peripheral blood primarily in adult populations, leaving uncharacterized tissue-resident lymphocytes collected from children. Here, we paired genotyping with gene expression profiling across four populations of tonsil-derived immune cell types – including previously uncharacterized germinal center B cells – collected from 103 children across development (ages 1-18). Using these data, we report cell-type specific gene networks and identify 13,393 eGenes (1,793 eGenes not previously reported in similar datasets) influenced by 27,603 eQTLs (5,199 not previously reported). We link discovered eQTLs to associations identified in pediatric and adult asthma and atopy traits, nominating 78 eGenes like *TRAF3*, *ZBTB10* and *JAZF1* in disease relevant cell types. Our resource is freely available and exemplifies the importance of discovery in native tissues and across human development.

## Introduction

Atopic inflammation preferentially affects barrier tissues—skin, gut, nasal mucosa, and lungs—that are exposed to environmental and dietary allergens. Central to the chronic and systemic nature of atopic disease is allergen-specific immunoglobulin E (IgE), which mediates symptoms and perpetuates disease. IgE class switching occurs in terminally differentiating antibody-secreting B cells within secondary lymphoid and mucosal tissues. The decision to class-switch to IgE, rather than another non-atopic isotype, is informed by B cell sensing of complex signals from specialized T helper cells and the tissue microenvironment^1–3^. Once established early in life, a predilection towards IgE production propels serial progression of anatomically diverse IgE-mediated diseases across the lifespan, including atopic dermatitis, food allergy, allergic rhinitis, and allergic asthma^4^.

Although allergen exposure is required to elicit allergic symptoms, genetic predisposition is a foundational contributor to atopic disease^5^. Nearly two decades of genome-wide association studies (GWAS) have identified thousands of single-nucleotide polymorphisms (SNPs) enriched among allergic subjects. Because most of these variants lie in non-coding regions, their target genes, the direction of transcriptional effects, and relevant cell types remain largely unresolved. One approach to infer such relationships is the integration of genotype and RNA-sequencing data to define expression quantitative trait loci (eQTLs) in disease-relevant tissues. The large eQTLGen study ^6^, which analyzed over 30,000 whole-blood samples, necessarily conflated diverse circulating cell types. The Database of Immune Cell eQTLs/Expression/Epigenomics (DICE) ^7,8^ improved cellular resolution by profiling 13 circulating immune cell types. However, tissue-resident, follicular, and mucosal lymphocytes—critical for IgE generation—were not included. Moreover, both resources only enrolled adult subjects, limiting relevance to pediatric-onset atopy, while we know the immune system changes across development^9–11^. Although additional single-cell resources are anticipated, they are not yet publicly available ^12,13^.

Given limitations of available data, many genome wide associations with atopy remain without an obvious connection to gene or immunologic mechanism. To address this gap, we collected excised tonsils—a mucosal, secondary lymphoid tissue—from 103 immunocompetent children. We generated eQTLs from naïve B cells, germinal center (GC) B cells, naïve T cells, and T follicular helper (Tfh) cells, and further annotated these data with ATAC-seq–defined open chromatin and Capture-C chromatin looping (**Figure 1**). We then statistically aligned associations with pediatric atopy-related GWAS signals to tonsillar eQTL signals we discovered here to nominate candidate effector genes and cell types for the GWAS traits. The strongest associations were observed for pediatric asthma, for which we confirm 16 previously implicated genes and identify 20 additional candidates. This resource provides a pediatric, tissue-relevant framework for linking genetic risk to transcriptional regulation in naïve and activated immune cells, with direct relevance to childhood atopic, inflammatory, and infectious disease.

**Figure 1:**
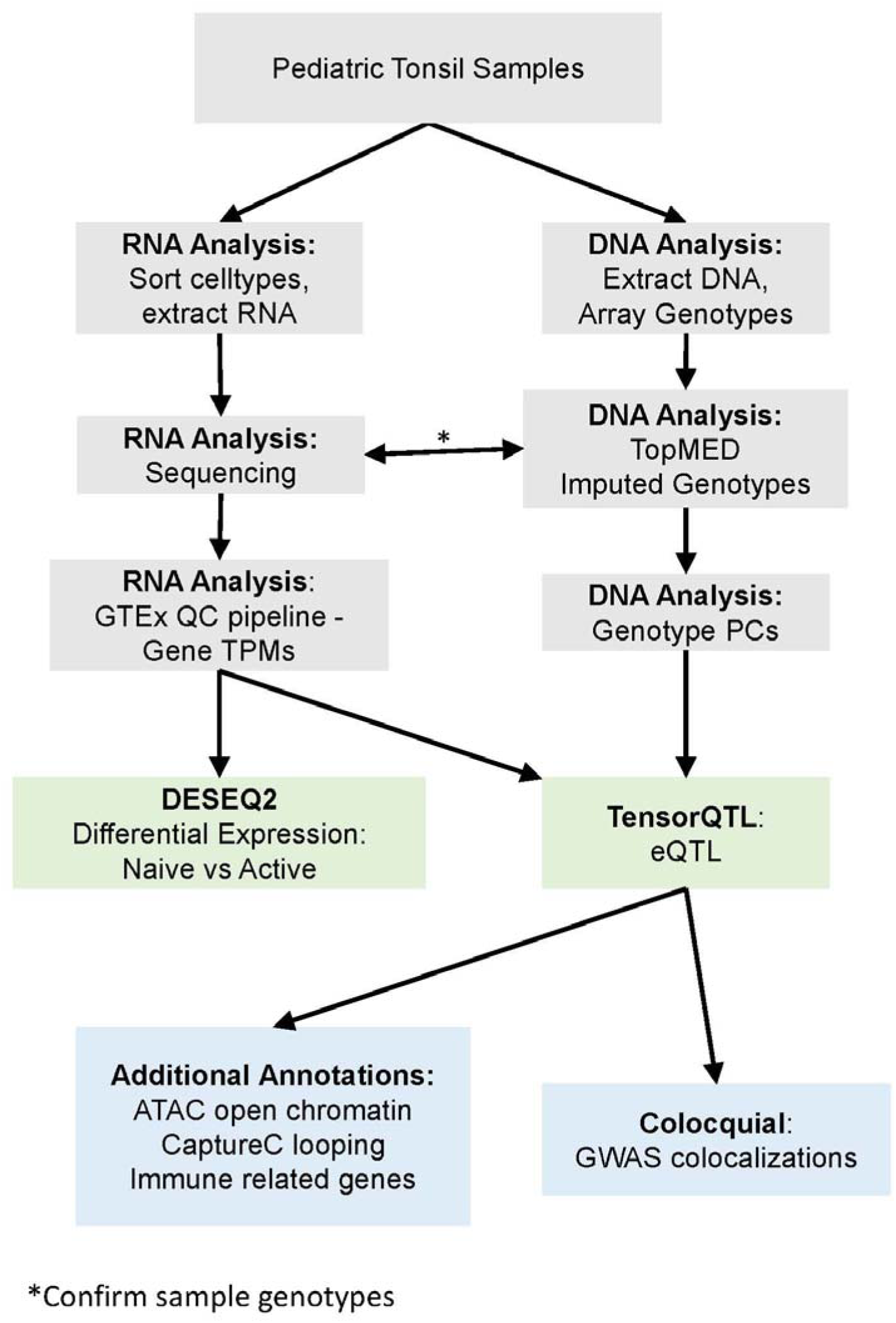
Analysis Overview Flowchart.

## Results

### Sample Collection and Information

Surgically excised, discarded tonsil pairs were collected from immune competent subjects ranging in age from 1-18 (average age 6.2 years). 56% of subjects were male. Indications for tonsil removal were not exclusive and included obstructive disorders (80% of subjects) such as sleep disordered breathing and sleep apnea, infectious disorders (15% of subjects) such as recurrent tonsillitis, and unknown diagnosis (7% of subjects).

### Pediatric Tonsillar Lymphocyte eQTLs

To identify cis-eQTLs, we assessed variants within ±1MB of each gene’s transcription start site (TSS) in our four pediatric cell types using TensorQTL^14^ (**Methods**). We found a total of 13,393 genes with at least one significant SNP in at least one cell type (eGenes), with between 6,594 and 7,145 eGenes per cell type (**Supplementary Table 1**). Of these eGenes, 5,905 (44%) are identified in a single cell type, 2,367 (17.7%) are identified in all four cell types, 1,587 (11.8%) in either only B or T cell types, and the remainder in mixed sets of 2 or 3 cell types (**Figure 2A**).

**Figure 2:**
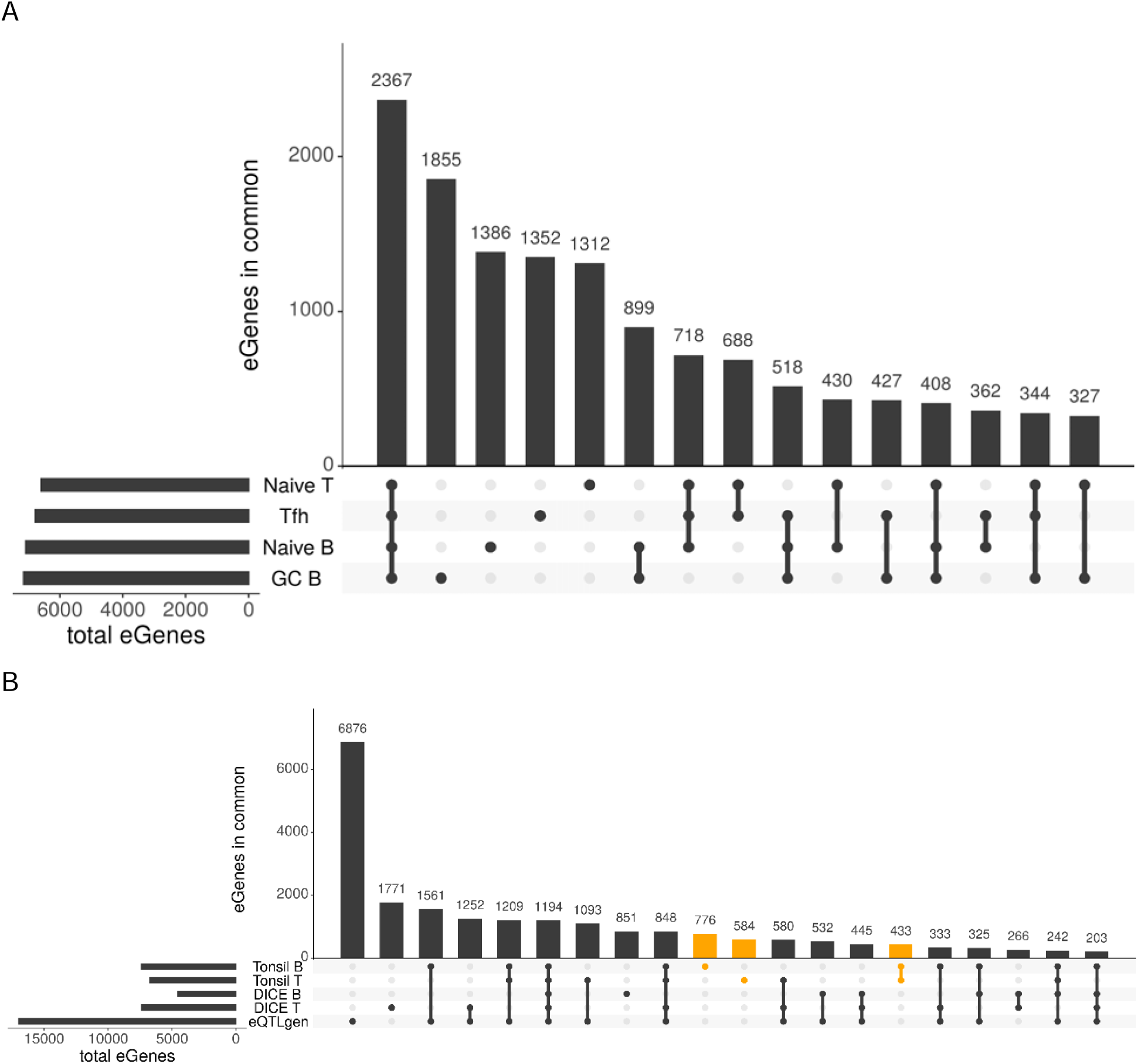
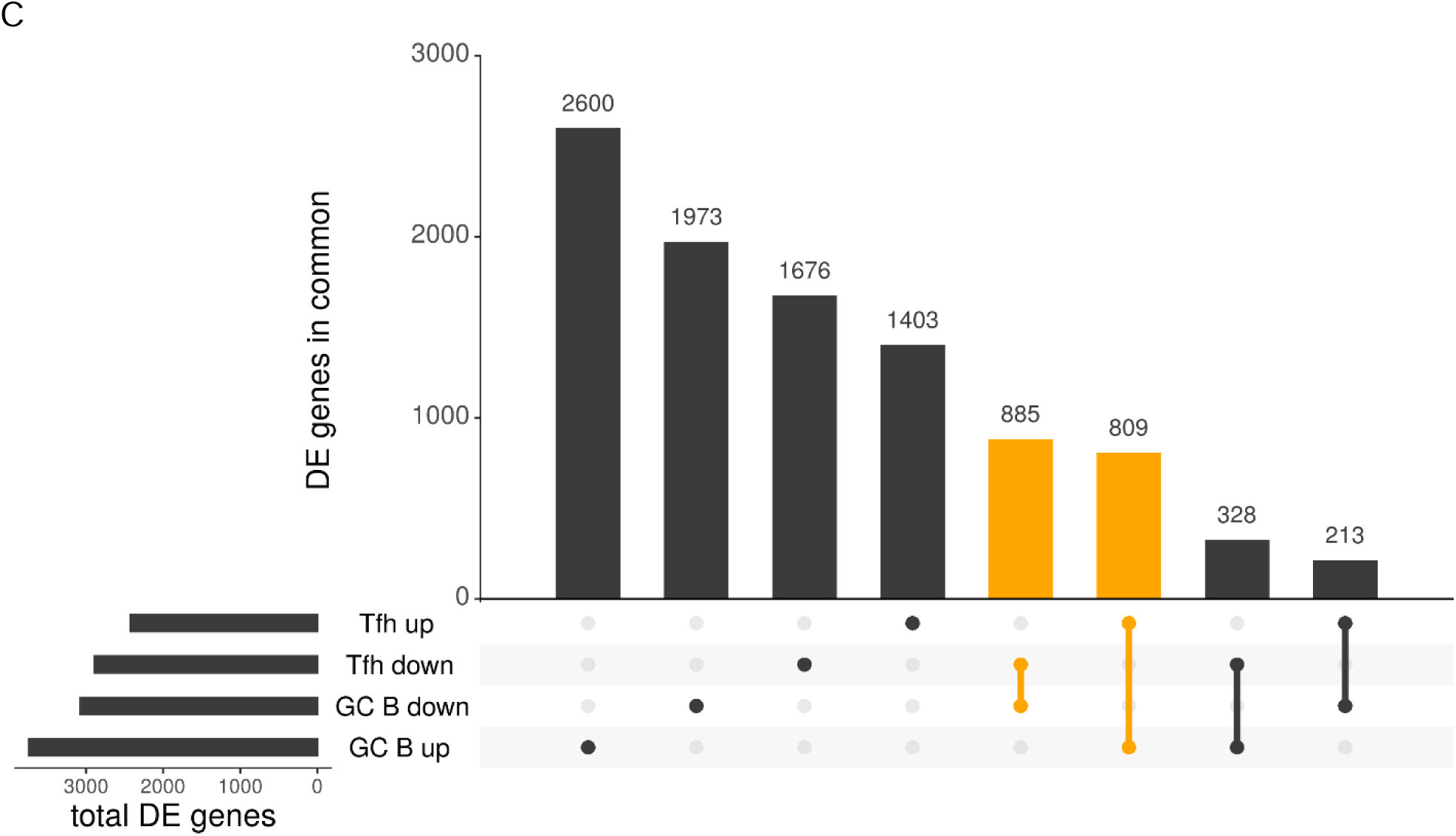
UpSet plots comparing differentially expressed genes and eGenes. UpSet plots showing the overlap between significant genes. A) Comparing all eGenes identified in our samples, across naïve B, GC B, naïve T, and Tfh cell types. B) Comparing all genes tested in all three studies, with eGenes identified uniquely by our samples (Tonsil B and Tonsil T) highlighted in orange. Shows only the top 20 intersections (31 total) C) Differentially expressed (DE) genes identified by DESEQ2 in naïve B vs GC B and naïve T vs Tfh are shown, divided into genes which have increased(up) or decreased(down) expression in the activated cell type. Genes which have the same direction of expression in the activated cell type are highlighted in orange.

To assess the validity of discovered eQTL, we referenced our discoveries against several databases of known immune genes and SNPs. First, we found enrichment for genes present in the Panther^15^ immune system gene set: 269/586 pathway genes are eGenes (46%, permutation p value = 0.01). In addition, we identified enrichment in relevant cell types for genes previously identified in B^16^ but not T^17^ cells in Gene Set Enrichment Analysis (GSEA) datasets: 389/764 B cell genes are eGenes (51%, permutation p-value < 0.0001) but 152/372 T cell genes are eGenes (41%, permutation p-value = 0.17), possibly due to the GSEA B cells better matching the GC cell subtype. On a SNP level, we considered overlaps between tonsil eQTL and SNPs identified in a study of pediatric-onset autoimmune diseases (pAIDs). We identified 44 eGenes with a lead SNP or proxy (**Methods**) identified in the pAID analysis, suggesting possible gene links and tissues of action for these variants. These data confirm that our eQTL analysis identifies immune relevant genes (eGenes for each above category are listed in **Supplementary Table 1**). We expect that some of these common expression variations may contribute to complex atopy traits via known immunologic pathways.

We then confirmed the physical connections between our 27,603 eQTLs and their eGenes. We found 6,926 eQTLs (25%) have a significant SNP within the 1,000 bp upstream of their eGene’s TSS, leading us to categorize them as likely promoter resident eQTL. Of the more distal eQTLs, 6,884 eQTLs (25%) have a cell type matched promoter-focused Capture-C loop^18,19^ between a significant SNP and the eGene. Finally, we found that most eQTLs had a significant SNP in an open chromatin region^18,19^ (i.e., ATAC-Seq peak) in the same cell type: 18,257 eQTLs (66%), as might be expected for regulatory variants.

To place these eQTLs in a larger context of previous discovery, we compared to those identified from a large-scale study from whole blood (eQTLgen; n=31,684 individuals) as well as an immune based dataset (DICE; naïve B, naïve T, and Tfh cells; n=91 individuals). Across our 4 cell types and eQTLs we identified, 5,199 of our eQTLs (19%) were not previously reported in either DICE or eQTLgen. On immediate inspection, we noted that we identified many more eGenes per cell type than the DICE datasets (we found 6,500+ eGenes per cell type; DICE found at most 3590 eGenes per cell type). We suspect this is due to differences/improvements in imputation reference panels (DICE: 1000G imputation, whereas we used TOPMed v4, **Methods**). To provide a fair comparison metric, we exclude from this comparison any eGenes where the lead variant or proxy (**Methods**) was not tested in the comparison dataset. After excluding those genes, we identified 2,781 eGenes not significant in the eQTLgen whole blood dataset, as well as 5,445 and 3,852 eGenes not present in the DICE adult B and T cell types, respectively. Comparing all together, we find 1,793 eGenes that were tested but not significant in either the eQTLgen or DICE datasets (**Figure 2B**). Interestingly, even though the GC B cell type was not tested in DICE, a full 1,017 of the novel eGenes are from the T cell types, suggesting the novelty of this dataset is not solely explained by the previously uncharacterized cell type. We hypothesize that these novel eGenes are enriched for either pediatric-involved biology (compared to adults) or tissue-embedded immune biology (compared to circulating immune cells), as these are the overlapping differences between our dataset and the DICE eQTL.

We next performed a series of age and sex stratified analyses (**Methods**) to identify eQTLs which are associated only with boys, girls, younger (≤ 4 years) or older (≥ 5 years) individuals in our dataset. While a direct interaction analysis performed on the pooled eQTL data did not yield any significant results after multiple hypothesis correction (**Supplementary Table 2**), when we split each cell type into subsets and called eQTLs, we identified 40-242 eGenes in one sex but not the full dataset and 81-143 in one age group but not the full dataset (**Supplementary Figures 1 & 2** and **Supplementary Table 3**).

### Colocalization with Asthma and Atopy Traits

To link these immune cell eQTL with specific atopy traits, we selected GWAS for asthma^20^, pediatric asthma^20^, atopic asthma^21^, child asthma^21^, atopic dermatitis/eczema (ADE)^22^, and a combined asthma, eczema or allergy phenotype (hereafter referred to as “allergic”)^23^ (**Supplementary Table 4**). Since we only colocalized regions with both a significant GWAS signal and a significant eQTL, we used a colocalization threshold of 0.8 conditional PP4 (**Methods**). By this metric, we identified a total of 200 colocalizations (trait-cell type-gene pairs) involving 78 unique genes across 6 traits (**Supplementary Table 5**). To identify a collection of variants statistically likely to harbor the causal variant, we tabulated approximate Bayes factor (ABF) credible sets for all significant eQTL (**Supplementary Table 6**).

We next sought to interpret these colocalization in the context of previous eQTL data sets. We confirmed that all our colocalizing lead eQTL SNPs were tested in the corresponding DICE cell type or eQTLgen. However, 141 colocalizations involve an eQTL not found in the corresponding DICE cell type and 34 were not found in eQTLgen. Of our 78 colocalizing eGenes, 25 also colocalized in DICE, and 53 were novel to our data (**Supplementary Table 7**). Several colocalizations involved genes that were high likelihood candidate genes for atopic GWAS: *ZPBP2*, *ORMDL3*, *IL6R*, and *TLR1*. While the eQTL in *ZPBP2* (naïve B) and *ORMDL3* (naïve T, Tfh) are present in DICE and colocalize, those in *IL6R* and *TLR1* are unique to our dataset and represent known biology we validate. *IL6R* is a primary immunodeficiency gene, loss of which is known to result in a syndrome that includes eczema symptoms,^24^ consistent with our colocalizing an *IL6R* eQTL in naïve T cells with ADE. Similarly, *TLR1* expression levels have previously been implicated in asthma pathogenesis^25^, so it is reassuring that we identify a colocalization between a Tfh *TLR1* eQTL and childhood asthma.

Of our 200 atopy trait colocalizations, 59 (29.5%) have a significant eQTL SNP that overlaps the gene promoter (1,000 bp upstream of the TSS), suggesting a mechanism of action in a specific immune cell type or types. As remaining colocalizations appear to be distal in nature, we next investigated overlap between our QTL loci and two additional previously published data types ^18,19^: cell type matched open chromatin (obtained via ATAC-Seq) paired with chromatin looping data (obtained from promoter Capture-C). While 10 (0.05%) have no overlaps with cell type matched open chromatin or loops, 129 (64.5%) colocalizing eQTL have at least one significant SNP that overlaps open chromatin and 49 (24.5%) overlap a matching loop to their eGene; 47 (23.5%) colocalizing eQTL have both. These 47 colocalizations span 21 genes and all 6 tested traits, including well known immunity genes like *AP1S3*, *TRAF3*, *RUNX3* and *SMAD4*.

We focus on pediatric asthma as an exemplar for the utility of matching eQTL source (pediatric, tissue resident) to trait, as the previous GWAS paper included a list of likely targets based on linkage but not formal colocalization analysis^20^. Of their 93 proposed genes for pediatric asthma, our colocalization analysis validates 16 of these as eGenes, including *TRAF3*, *JAZF1*, *TNFSF4*, and *OVOL1*. Only 7 of these eGenes were found to colocalize in DICE data. In addition, we find 20 non-nominated eGenes colocalizing with pediatric asthma, only 4 of which are found by DICE (**Supplementary Table 8**). These eGenes include several excellent candidate genes, including *EEFSEC*, *ZBTB10*, and *TNFSF11*. *EEFSEC* is a particularly interesting candidate, given its role in selenocysteine incorporation^26^, links to COPD^27^, and the history of controversy regarding selenium levels affecting asthma^28^.

We next turned to identify relevant cell types across our colocalized eQTL variation. For 62 eGenes (78.5%) corresponding to 89 eGene-trait pairs, the colocalizing target is an eGene across multiple tested cell types. We narrowed these to 32 eGenes (49 eGene-trait pairs) with overlapping looping data, and then compared the between cell type eQTL, requiring a conditional PP3 greater than 0.8 to call them distinct signals. By this metric, 13 eGenes (21 eGene-trait pairs) show cell type specific regulation, with distinct eQTL in multiple cell types supported by looping data.

A leading, aligned example of cell type resolution emerged from the known immune activation gene, *TRAF3*. This region harbors a loop between our eQTL signal in GC B cells in the promoter (red line denotes top SNP), while the naïve T eQTL is within the gene body (black line denotes top SNP, **Figure 3**). Both eQTL colocalize with the ADE trait, while only the naïve T eQTL colocalizes with pediatric asthma (**Figure 3**), suggesting that the differential regulation might relate to different facets of atopy traits at this locus. Fine mapping helps to further narrow a subset of SNPs likely to be causal for each cell type specific eQTL (i.e., gray highlights identify the naïve T eQTL credible set).

**Figure 3:**
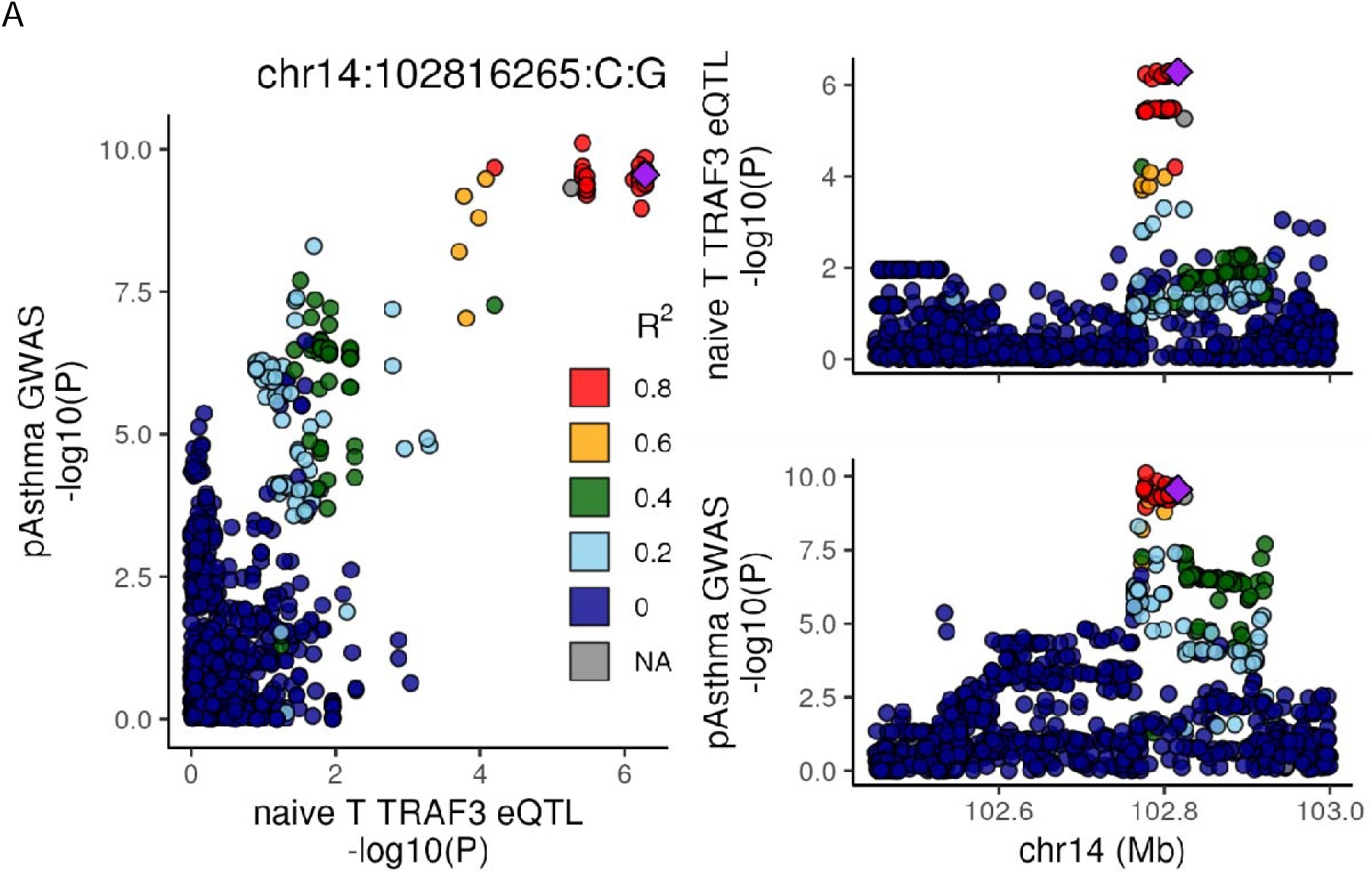
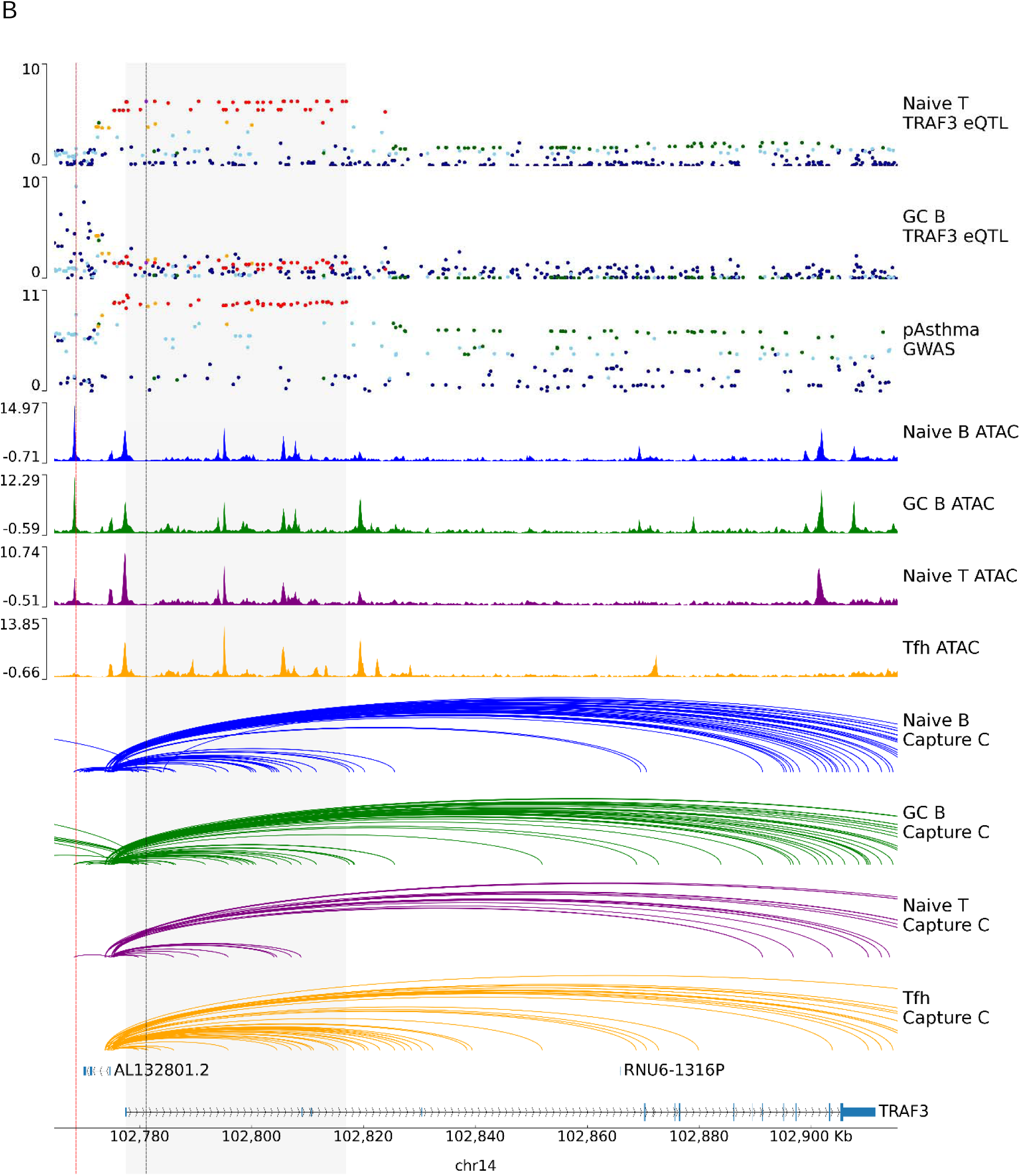
Colocalization, looping and ATAC peaks around TRAF3. A) LocusCompare plot for colocalization between pediatric asthma and naïve T TRAF3 eQTL. SNPs in all plots are colored according to LD to chr14:102816265. B) The first 3 tracks show the locus plots for naïve T TRAF3 eQTL, GC B TRAF3 eQTL, and pediatric asthma GWAS. Naïve B ATAC and Capture C loops shown in blue, GC B in green, naïve T in purple, and Tfh in orange. Vertical black line indicates naïve T TRAF3 eQTL lead SNP; vertical red line indicates GC B TRAF3 eQTL lead SNP; vertical gray highlight indicates TRAF3 naïve T eQTL ABF credible set. The TRAF3 GC B eQTL credible set is a single SNP and corresponds to the vertical red line.

In addition, we observed that *ZBTB10* colocalized with 5 of our 6 tested traits (excluding ADE) only in the Tfh cell type, with significant SNPs overlapping both open chromatin and loops to the *ZBTB10* promoter (**Figure 4**). *ZBTB10* is a gene often linked to asthma and atopy traits without a clear mechanism^29,30^. While *ZBTB10* was initially identified in an eQTL study of bronchial tissue^31^, here we connect this locus and gene to a mechanism of action in Tfh cells.

**Figure 4:**
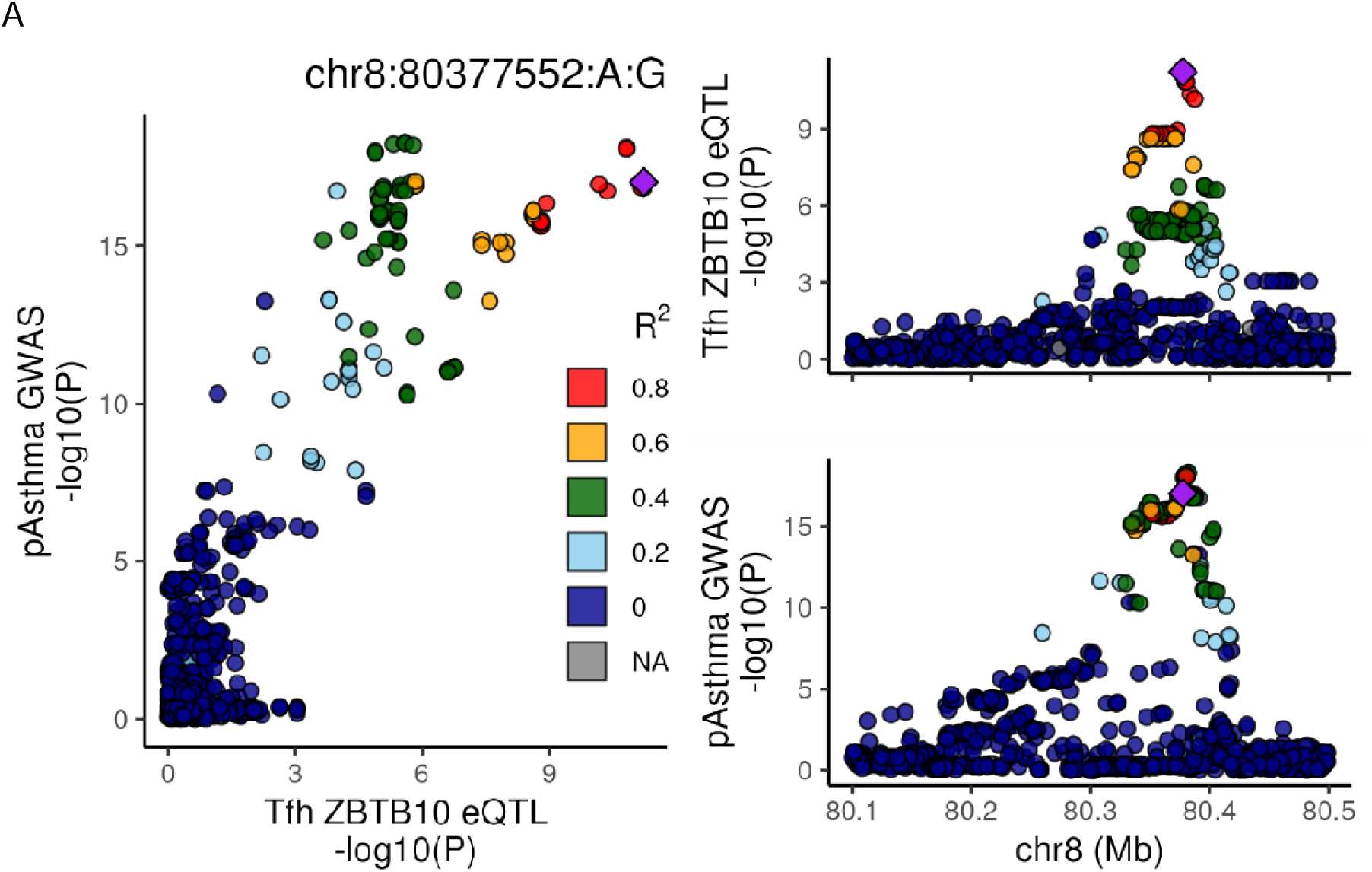
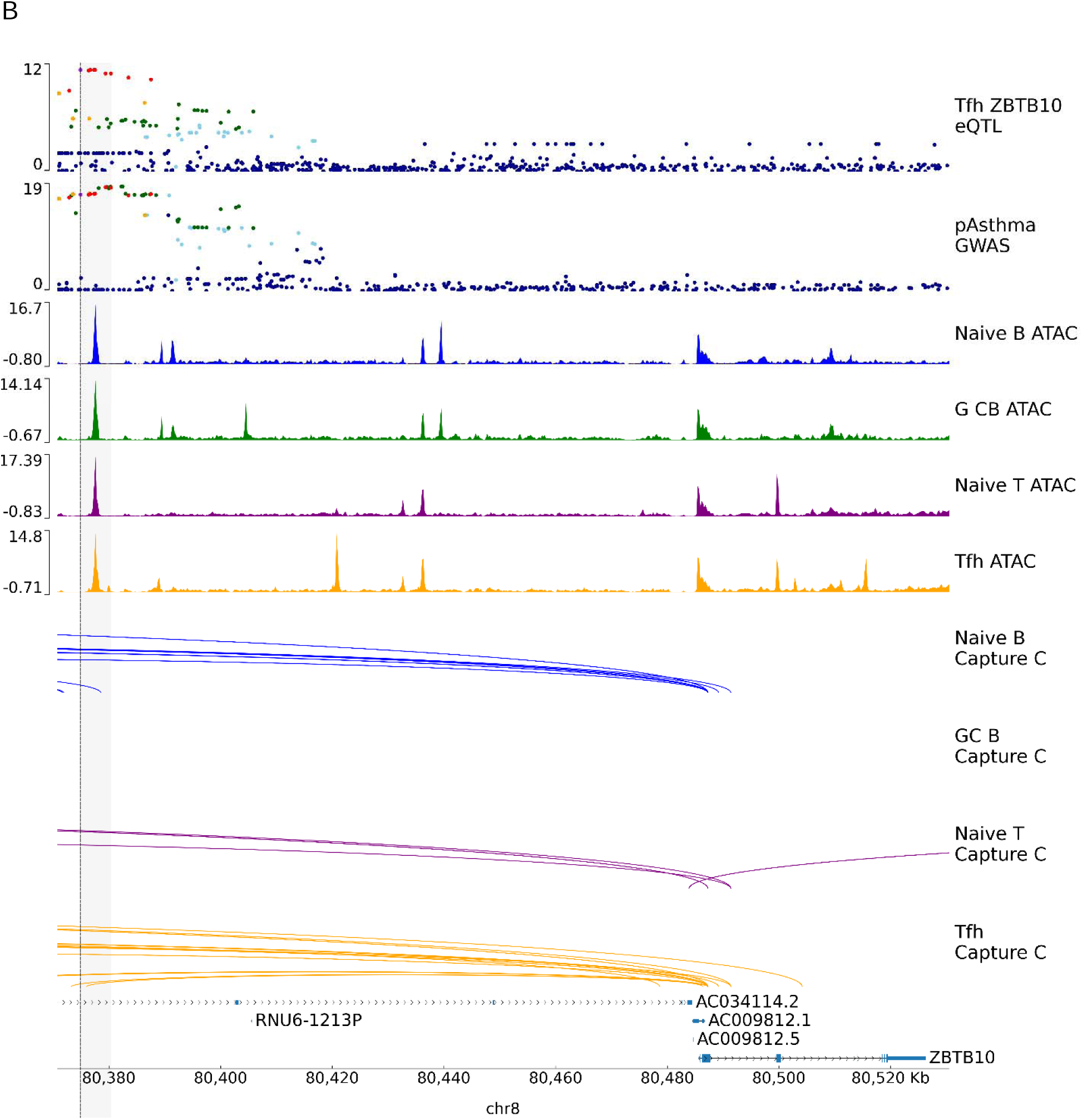
Colocalization, looping and ATAC peaks around ZBTB10. A) LocusCompare plot for colocalization between pediatric asthma and Tfh ZBTB10 eQTL. SNPs in all locuszoom plots are colored according to LD to chr8:80377552. B) The first 2 tracks show the locus plots for Tfh ZBTB10 eQTL, and pediatric asthma GWAS. Naïve B ATAC and CaptureC loops shown in blue, GC B in green, naïve T in purple, and Tfh in orange. Vertical black line indicates Tfh ZBTB10 eQTL lead SNP; Vertical grey highlight indicates it’s ABF credible set.

A final example of focus involves *JAZF1*, a significant eGene across all four tested cell types. However, *JAZF1* colocalizes with pediatric asthma (**Figure 5A**) and combined allergic traits in both naïve B and naïve T cell types, but not in either differentiated cell types. Assessing the conditional PP3 and PP4, we find the naïve signals colocalize with each other (conditional PP4 = 1, **Figure 5B**) but are distinct from their respective more differentiated counterpart eQTL signals (conditional PP3 = 0.92 for B cell comparison and PP3 = 1 for T cells; **Figures 5C and 5D**). This suggests a mechanism of action for atopy traits that occurs in the naïve B and T cell types, prior to more terminal differentiation. Further, the direction of effect aligns with the known biology of *JAZF1*, where an increase in expression is associated with a decrease in inflammation^32,33^ as well as lower atopy risk.

**Figure 5:**
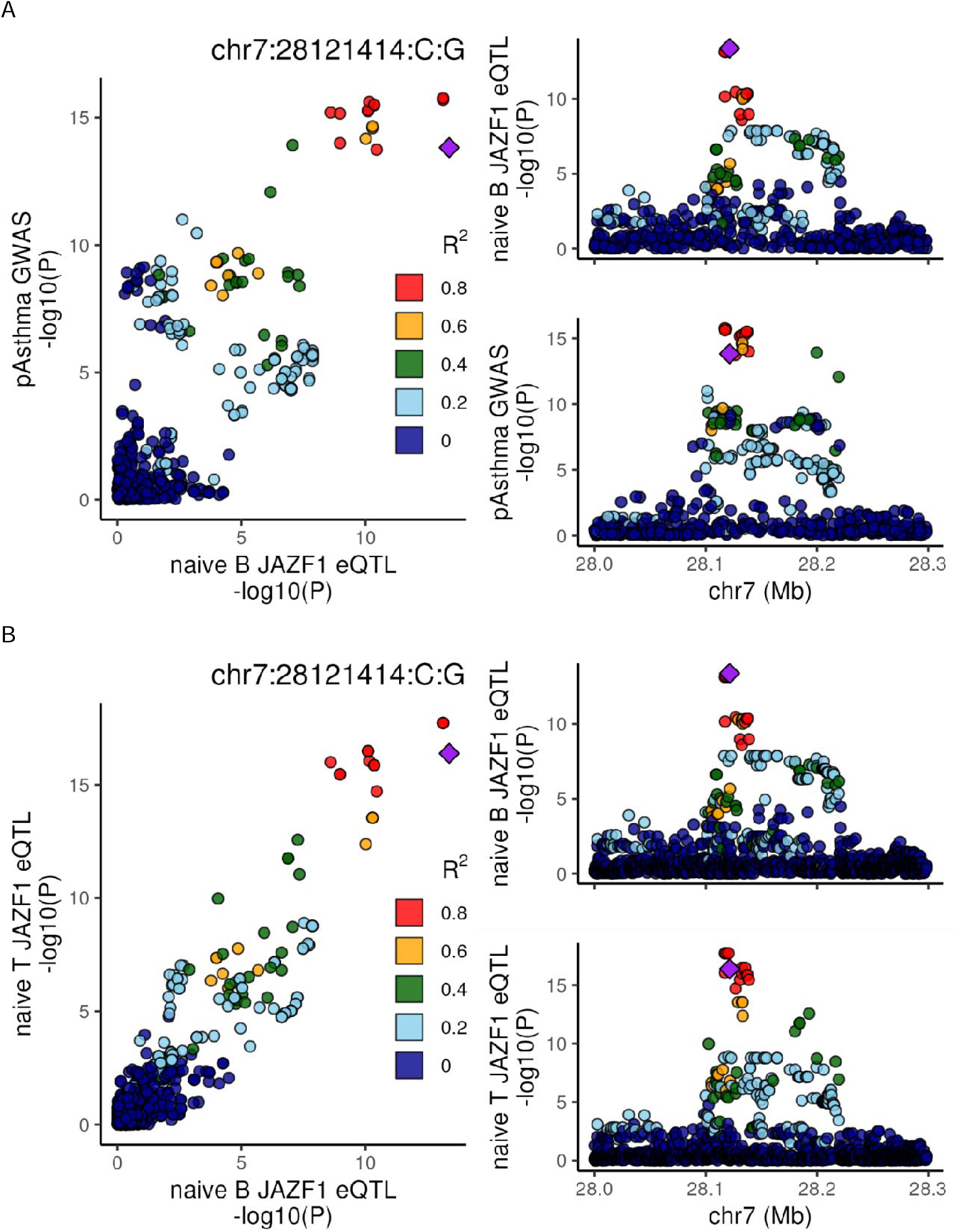
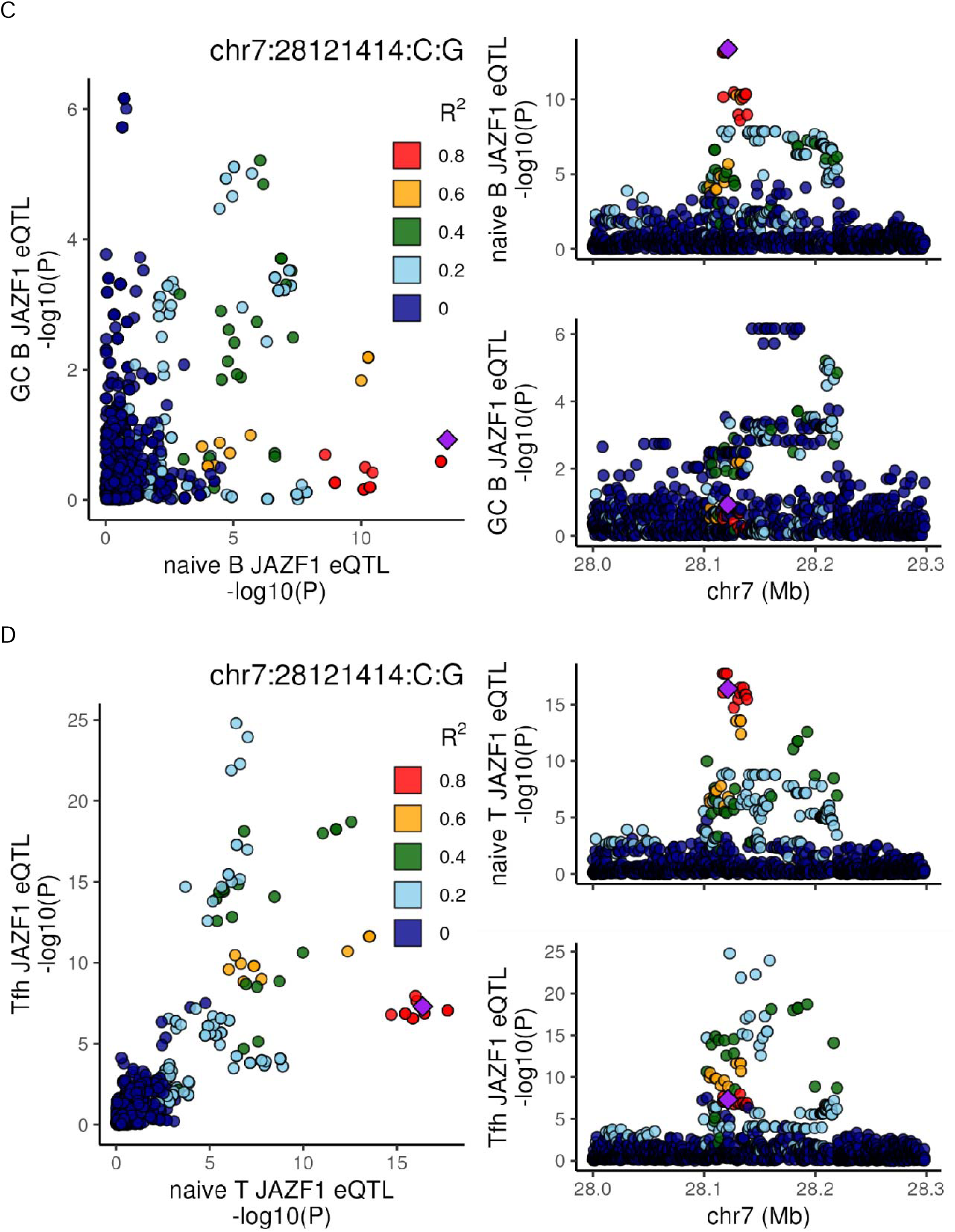
JAZF1 Colocalizations. LocusCompare plots demonstrating colocalizations between A) pediatric asthma and JAZF1 expression in naïve B cells, B) JAZF1 expression in naïve B and T cells. And non colocalizations between C) JAZF1 expression in naïve B and GC B cells, and D) JAZF1 expression in naïve T and Tfh cells. In all cases LD information used from our eQTL samples, SNPs are colored by their LD to the lead naïve B eQTL SNP, and genomic positions are in hg38.

### Pediatric Tonsillar Lymphocyte Transcriptomes

Finally, we assessed differentially expressed genes across our samples. After identifying genes expressed in each cell type (**Methods**), we considered only genes with at least 10 reads across all samples, we consider as differentially expressed a gene with a Bonferroni corrected significant log fold change of ≤ -1.5 or ≥ 1.5. By these criteria, we find 5,314 differentially expressed genes between naïve T and Tfh cells, 6,808 between naïve B and GC B cells, 9,588 between GC B and Tfh cells, and 6,722 between naïve T and naïve B cells (**Supplementary Table 9**, **Supplementary Figure 3**). Comparing naïve vs. more differentiated cells, 2,235 genes are identified as differentially expressed in both B and T cell types, with 1694 (75.8%) being directionally concordant (**Figure 2C**).

We also compared naïve vs. more differentiated subsets across metadata variables: males, females, younger (≤4 years of age) and older (≥ 5 years of age) samples, resulting in slightly lower but similar numbers of differentially expressed genes (**Supplementary Table 10**). We identify 3-50 heterogeneous genes of interest across age and sex subset pairs (**Methods**, **Supplementary Figures 4 & 5**). While some slight differences in expression due to age or sex are identified, no asthma or atopy related genes are identified.

Extending the differential expression to the eQTL analysis, we find that more than 1,200 eGenes per cell type were differentially expressed between the relevant naïve and more differentiated lymphocytes; these genes are likely to be involved in B or T cell interaction, activation and trafficking to the follicle. (**Supplementary Table 1**). Within the pool of 79 atopy GWAS colocalizing genes, we find 18 differentially expressed genes between naïve T and Tfh cells and 12 between naïve B and GC B cells (26 genes total). These genes include *JAZF1*, *RUNX3* and *ZBTB10* in T cells, consistent with their eQTL effects.

## Discussion

In this study, we generated a catalog of pediatric immune cell type-specific eQTLs from tonsil-derived lymphocytes across four cell populations: naïve B cells, GC B cells, naïve T cells, and Tfh cells. From 103 children (all <18), we identified 27,603 eQTLs, with 5,199 eQTLs representing novel discoveries not previously reported in existing databases. Importantly, 44% were identified in a single cell type, with 12% in either only B or T cell types, emphasizing the gains of profiling distinctive cell populations. Supporting the relevance of our identified eGenes, we found that there is significant enrichment for established B and T cell markers. Through colocalization analyses with pediatric and adult GWAS, we identified 78 candidate causal genes, confirmed 16 previously proposed candidate genes for pediatric asthma and nominated 20 additional gene candidates, demonstrating the value of this resource for mechanistic insights into immune-mediated diseases.

Our findings help to fill a critical gap in the existing eQTL landscape and complement available resources. Prior large-scale efforts, including DICE^7^ and eQTLgen^6^, have focused predominantly on adult-derived peripheral blood samples, which may not fully capture the developmental dynamics and cell type specific regulatory architecture relevant to pediatric disease. By profiling tissue-resident lymphocytes across childhood development, our resource provides new insights into gene regulatory networks that may be particularly relevant during critical windows of immune system maturation. The inclusion of GC B cells—a population not comprehensively characterized in previous eQTL studies, as it is not present in circulating blood—is especially significant, as these cells play central roles in antibody maturation and are dysregulated in multiple inflammatory conditions^34–37^. Our data thus enables investigation of developmental and tissue-specific regulatory mechanisms that would be missed by studies restricted to adult peripheral blood.

Several of the novel candidate genes we identified through colocalization with asthma and atopy GWAS signals have compelling biological connections to immune function and airway inflammation. For instance, TRAF3 plays critical roles in B and T cells relevant to allergic disease. Through a STAT6 mechanism, TRAF3 promotes skewing of naïve helper cells into IL4-secreting Th2 cells that promote IgE class-switching,^38^ while in B cells, TRAF3 restrains class-switching to IgE by interfering with CD40 signaling^39^. *ZBTB10*, which we nominated as a candidate gene for asthma in Tfh cells, is a transcription factor known to regulate immune cell differentiation and has been implicated in telomere biology^30,40^—processes especially relevant to activated T cells, which are known to reactivate telomerase to become effectively immortal^41^. Similarly, other genes identified in our analyses participate in B cell receptor signaling^42^, T cell activation^43–46^, cytokine production^47,48^, and regulation of inflammatory responses^32,33,49^, all of which are central to IgE-mediated atopic diseases including allergic asthma. The cell types implicated by our eQTLs further suggest that genetic risk for atopy may operate through distinct mechanisms in different immune cell populations. For example, eQTLs specific to Tfh cells may influence the balance of helper T cell responses that drive B cells to class-switch to IgE^50,51^, while GC B-specific eQTLs could promote choosing IgE, over IgG or IgA, due to altered sensitivity to T cell help and/or inflammatory mucosal signals^52–54^. Beyond asthma, the genetic architecture we uncovered is relevant to other atopic conditions including allergic rhinitis, eczema, and food allergies, as these traits share common genetic underpinnings and immunologic mechanisms. Our resource can therefore facilitate discovery across the spectrum of pediatric allergic and autoimmune diseases, helping to elucidate shared versus disease-specific regulatory mechanisms.

Our study carries several limitations that are worth acknowledging. First, our analyses were restricted to tonsillar lymphocyte datasets. Although the adaptive immune response is a critical perpetuator of atopic diseases, numerous other hematopoietic and non-hematopoietic cell types also contribute. For this reason, it is not surprising that established atopy-associated genes expressed in myeloid cells (e.g., *FCER1A)*, keratinocytes (e.g., *TSLP* and *FLG*) and airway smooth muscle cells (e.g., *ADAM33*), but not tonsillar lymphocytes, were not implicated by our study. We anticipate that similar analyses of eQTL datasets generated from non-lymphoid cell types would implicate both known and novel egenes. Second, we did not include adult samples in our study. This design choice was deliberate, as our goal was to characterize the pediatric immune landscape, but it means we cannot fully disentangle whether the discoveries made here but not observed in adult eQTL resources arise from developmental stage versus tissue of origin. Third, while our sample size is large from the point of view of pediatric collections, it is modest compared to very large blood-based eQTL studies with thousands of participants. This limits statistical power to detect eQTLs with small effect sizes, or lower frequency variants with possibly stronger effects, although notably our strategy of discriminating between tissue-resident lymphocytes in 103 tonsil donors yielded almost as many eGenes as eQTLgen did with only 0.35% as many subjects. Fourth, while we did not collect ancestry data on these samples we can infer from the principal component analysis that they are largely European-like (56%), with only a few Asian- and African-like patients. A more ancestry-inclusive analysis would likely find additional signals tagged by variation common in those groups, but rare or absent in European-like groups, though it would require similarly inclusive trait GWAS. Finally, tonsils were collected as part of routine clinical care for either infectious or obstructive indications, and we cannot rule out the possibility that the underlying clinical condition influenced gene expression patterns. However, we did not observe systematic differences between diagnostic groups in our quality control analyses, and our successful replication of known immune genes suggests that any such effects are likely modest.

This work represents an important step forward in understanding the genetic architecture of pediatric immune function and immune-mediated disease. By providing the first comprehensive catalog of eQTLs in tissue-resident pediatric immune cells, including the previously uncharacterized GC B population, we enable mechanistic interpretation of genetic associations for asthma, atopy, and related autoimmune conditions in children. Our resource is publicly available and will serve as a foundation for future studies investigating the interplay between development, tissue context, and genetic regulation in pediatric immunology.

## Methods

### Tonsillar T and B cell subset preparation and sorting

Fresh tonsils were obtained as discarded surgical waste from 105 de-identified immune-competent children undergoing tonsillectomy to address airway obstruction or recurrent tonsillitis. The average age of tonsil donors was 6.2 years, 56% were male. Since they were de-identified, analysis of collected tonsillar tissue was determined to not be human subjects research by the Children’s Hospital of Philadelphia Institutional Review Board. A single cell suspension of tonsillar mononuclear cells (MNCs) was created by mechanical disruption (tonsils were minced and pressed through a 70-micron cell screen) followed by Ficoll-Paque PLUS density gradient centrifugation (GE Healthcare Life Sciences).

MNCs from each sample were split into three fractions. One fraction was set aside for DNA extraction. The second fraction was depleted of CD19-positive cells (StemCell) and enriched in CD4+ T cells (Biolegend) with magnetic beads before sort separating viable naïve T cell (CD4^+^CXCR5^-^PD1^-^CD45RO^-^) and Tfh cell (CD4^+^CD45RO^+^CXCR5^+^PD1^hi^) subsets. The third fraction was stained and immediately sort-separated into viable naïve B cells (CD19^+^CD21^+^CD27^-^IgD^+^CD38^-^) and germinal center B cells (CD19^+^CD21^+^IgD^-^CD38^+^) on a MoFlo AstriosTM (Beckman). Each sorted subset contained approximately 1×10^6^ cells and LIVE/DEAD Aqua (Thermo Fisher) counterstaining was used to ensure viability of sorted cells. T cell and B cell sorting strategies were performed as previously^18,19^(**Supplementary Figure 6**).

### DNA extraction and genotyping

Chromosomal DNA was extracted from 105 tonsil DNA samples using QIAamp DNA Blood Mini Kits (Qiagen). DNA quantity and quality was verified using a Nanodrop 2000 (Thermo Fisher). Genotyping was performed by the Center for Applied Genomics using Infinium Global Screening Arrays (Illumina).

We used Genome Studio 2.0.5^55^ to filter SNPs from the raw array calls if they had the following characteristics: AB T mean > 0.15, AB R mean < 0.3, call frequency < 0.5, cluster separation < 0.2748, het excess < -0.6 or > 0.4. Additionally, we removed all X chromosome SNPs with call frequency < 0.95. We filtered manually if the call frequency was between 0.5 and 0.75, het excess was between -0.6 and -0.35, or 0.2 and 0.4; removing SNPs in these zones without clearly clustered calls. We modified the recommended heterozygote excess filter due to our mixed ancestry population. Our final call set included array called genotypes for 566,408 SNPs.

We imputed genome wide variants using the TOPMED reference panel^56–58^ on GRCh38 for chromosomes 1-22 and X, with no allele frequency check and an R-squared filter of 0.3. TOPMED used 541,283 of our array genotyped SNPs for imputation. We subsequently analyzed imputed SNPs with MAF > 0.01 and imputation information score > 0.5, leaving 11,755,386 SNPs for downstream applications.

We calculated genotype principal components (PCs) and identity by decent (IBD) using plink v1.90b4.5^59,60^ (**Supplementary Figure 7**) and use 10 PCs as covariates when calling differentially expressed genes and eQTL. IBD analysis was performed on array called variants only, while PCs were calculated on imputed variants. After IBD analysis, we identified a trio of siblings (pihat ∼.5) and a pair and trio of cousins (pihat ∼ .125). We removed two of the three siblings from all analyses but due to small sample size retained the cousins. We further note that while we have no self-identified ancestry information, the PC analysis confirms roughly the expected population ancestry split, with 58 samples corresponding to a European-like grouping (56%), 26 to an African American-like grouping (25%), 7 to an Asian-like grouping (7%) and 12 of unclear ancestry (12%).

### RNA extraction, library preparation and sequencing

Immediately after sorting, each of the B and T cell subsets (i.e., 4 subsets of 105 subjects = 420 samples) were cryopreserved in TRIzol (Thermo Fisher) and stored at -80°C. Upon enrollment completion and before RNA extraction, samples were randomized into 24 groups of 18 samples per group to minimize potential confounding effects from batch. RNA was extracted and purified from sample groups on semi-consecutive days using Direct-zol RNA Microprep kits (Zymo Research). RNA quantity and integrity was verified with a 2100 Bioanalyzer (Agilent). 419 samples passed quality criteria (RIN score >6.5).

Libraries were generated using purified RNA samples normalized to 100 ng input with the NEXTFLEX® Rapid Directional RNA-Seq in the Sciclone G3 liquid handling instrument (following manufacturer recommendations). The method is designed to prepare directional, strand specific RNA libraries for sequencing using Illumina® sequencers. The procedure is ideal for insert sizes of >150 bp. Molecule directionality is retained by adding dUTP during the second strand synthesis step as only the cDNA strand is sequenced. This kit included the necessary reagents to process the user’s purified RNA sample through preparation and amplification for sequencing. The enriched RNA pool was fragmented, converted to cDNA, and PCR-ligated to indexes using an Illumina adaptor. The LabChip® GXII Touch HT was used to measure the molar concentrations of the indexed libraries and samples were pooled for sequencing. The concentration of each pool was also measured using an Illumina Library Quantification kit (KAPA Biosystems) and sequenced on an Illumina NovaSeq 6000 (Illumina, Inc, San Diego, CA) genome sequencer in two flow cells. A minimum of 40 million paired-end 100 bp reads per sample were obtained. After sequencing, sample randomization was broken for downstream analyses.

RNA sequencing processing was performed using the TOPMED/GTEx pipeline^61,62^ as a template. We used fastQC^63^ followed by multiQC^64^ to visualize raw sequencing statistics; all samples passed QC. Reads were aligned to GRCh38 using STAR 2.7.5c^65^ and we called variants in the aligned bam files using bcftools 1.9^66^. Comparing RNA-seq called variants with genotyped variants and genotyped sexes with expected sexes, we corrected mislabeled sample IDs. We marked duplicate reads using picard 2.23.3^67^ and calculated expression using RSEM 1.3^68^. We calculated read counts and tpm using RNA-seQC 2.3.6^69^.

### Reference files

For alignment, we used the hg38/GRCh38 reference genome^70^ downloaded 01/11/2022, including HLA but removing ALT and Decoy contigs. We used gene annotations from gencode v34^71^ downloaded 01/13/22. For LD lookups information, we used 1000 genomes phase 3 GRCh38 aligned data^72^, downloaded from the New York Genome Center on 07/25/2019. We annotated SNPs with rsIDs using dbSNP154^73^ downloaded from NCBI on 02/18/21.

### Additional covariates for modelling

In addition to the 10 genotype PCs already described, we calculated PEER^74^ covariates for each cell type and selected the number of covariates that maximized eGene number (12 PEER for naïve B, 11 for GC B, 13 for naïve T, 12 for Tfh). We included age, sex, diagnosis of infectious and diagnosis of obstructive as explicit covariates for eQTL analysis.

### Differential expression analysis

We identified differentially expressed genes using the DESeq2^75^ 1.40.2 R package. We performed a default paired analysis using a Wald test across four cell type pairings: naïve B vs GC B, naïve T vs Tfh, naïve T vs naïve B, and GC B vs Tfh, considering all genes with more than 10 reads. In addition to the full dataset, we also analyzed the data subsets male only, female only, younger (4 years old or younger) and older (5 years old or older) for the combinations naïve B vs GC B, and naïve T vs Tfh. To account for multiple analysis, we considered as differentially expressed genes with an adjusted p-value < 0.0025 (Bonferroni correction based on 20 tests performed - accounting for full dataset, male only, female only, younger, and older analyses across 4 cell type pairings), and a log_2_ fold change ≤ -1.5 or ≥ 1.5.

For the subset analyses, we tested for heterogeneity between sex and age groups using a Cochran’s Q test as implemented in the metafor^76^ R package. To reduce the multiple testing burden, we only tested for heterogeneity in autosomal genes with a log_2_ fold change difference in expression between the sex or age groups that was greater than 0.5. We considered the heterogeneity significant and of interest if Cochran’s Q test p-value (Qp) was less than Bonferroni correction based on the number of genes tested (range 4.99 × 10^-6^ to 8.84 × 10^-6^).

### eQTL analysis

We called eQTL using tensorQTL^14^, evaluating SNPs with a 1Mb window (+/- 500kb flanking) around the transcription start site (TSS) for each expression gene, including covariates in the analysis as discussed above. We considered significant genes where the gene q-val < 0.05 (calculated using the default ST procedure^77,78^) and a SNP in the region was below the permutation identified threshold. These criteria identified 7,088 eGenes in naïve B, 7,145 in GC B, 6,594 in naïve T and 6,776 in Tfh cells. UpSet^79,80^ plots were used to visualize overlapping gene sets.

To further assess the impact of age and sex on these eQTL, we performed 3 types of stratified analyses for each cell type. First, we used tensorQTL to directly model sex differences by effect modification, (i.e., Y ∼ SNP + COV + SEX + SNP*SEX or Y ∼ SNP + COV + AGE + SNP*AGE). Second, we split the data into separate pools and perform eQTL analysis in those strata, minus the relevant covariate. For sex, we stratified into male (total n = 58) and female (total n = 45); for age we stratified into two groups: aged four or younger (total n = 50) and five or older (total n = 53). These stratified eQTL were directly compared to the unstratified analysis to identify potentially novel eGene associations. Finally, we meta analyzed the stratified eQTL using a random effects model in METASOFT^81^ so that we could identify eGenes which have identifiable heterogeneity.

### eQTL colocalizations

We linked our catalog of eQTL (generated in this study or identified in the Database of Immune Cell e/QTLs/xpression/pigenomics (DICE) ^7^ study) with six atopy related GWAS traits using ColocQuial^82^ (a wrapper which automates application of pairwise statistical colocalization via coloc^83^): asthma^20^, atopic asthma^21^, child asthma^21^, atopic dermatitis/eczema (ADE)^22^, pediatric asthma^20^, and a combined asthma, eczema or allergy phenotype (allergic)^23^ (**Supplementary Table 4**). Summary statistics were lifted over to genome build hg38 prior to any analyses. We identified significant regions for colocalization by selecting reported lead variants with p-value < 5 × 10^-8^ and including flanking genomic regions until we obtained a 50kb flank without a variant with a p-value < 1 × 10^-5^. Regions used for colocalization for each trait can be found in **Supplementary Table 11** in the format used for ColocQuial submission. ColocQuial was run using default settings, and tested for colocalizations only if there was a significant eQTL (SNP < permutation threshold) in the given region. We therefore considered regions with PP4 ≥ 0.8 or PP4/(PP3+PP4) ≥ 0.8 as colocalized signals and regions with PP3 ≥ 0.8 or PP3/(PP3+PP4) ≥ 0.8 to be distinct signals, because all regions tested already have significant GWAS and eQTL signals.

### eQTL comparisons

To compare our eGenes with the existing DICE data^7^ available for naïve B, naïve T, and Tfh cells and whole blood eQTLgen data^6^, we first needed to identify SNP-gene pairs had been tested in both datasets, given differences between studies in imputation reference panels that were used: DICE and eQTLgen used 1000 Genomes Project as the reference data for imputation^6,7^, while we utilized TOPMed as our imputation reference panel. To identify novel SNP-gene pairs, we used 1000 Genomes Project (Phase 3) data to obtain proxy SNPs (LD r^2^ > 0.8 in EUR) for all lead variants identified in our datasets. If a lead or proxy was listed in the all pairs file for DICE or eQTLgen, we considered that signal we identified (and tagged by the SNP we identified) to have been tested in those data sets. For DICE, the assessment of novelty thus dropped 1665 eGenes for naïve T, 1720 for Tfh, 1836 for naïve B and 1774 for GC B celltypes from our data set. For comparisons to eQTLgen, we excluded from our own data 875 eGenes for naïve T, 868 for TFG, 918 for naïve B, and 885 for GC B. While we exclude them to be conservative in our estimation of novel eGenes, we acknowledge that given they were not identified previously, they are still new.

### eQTL overlaps

We assessed overlaps with additional, disease-relevant datasets on both an eGene and per SNP basis. For Panther^15^ gene set comparisons, we downloaded the list of human Immune System Process genes from https://www.pantherdb.org/ on 01/30/2025. We counted how many eGenes across our 4 cell types were present in this gene set and calculated a p-value based on 10,000 permutations of total genes tested in the eQTL analyses. Similarly, we downloaded GSEA gene lists relating to B cells^16^ (GSE12366 and GSE12845) and T cells^17^ (GSE40068) from https://www.gsea-msigdb.org/gsea/msigdb/human/genesets.jspon05/28/26. This analysis was done similarly to the above, but we considered only B cell (naïve B and GC B) tested genes for the B cell gene list, and likewise for the T cell gene list.

For SNP based comparisons to pediatric autoimmune diseases (pAIDs)^84^, we used summary statistics provided by the authors and considered an eGene positive if a SNP or LD r^2^ > 0.8 proxy is significant in the pAID meta-analysis. This study was analyzed in this fashion because it was genotyped using a targeted array, making it unsuitable for formal colocalization.

We considered a SNP in a gene’s promoter if it is within 1,000 bp of the TSS. We used bedtools^85^ intersect to look for overlaps with ATAC-seq data^18,19^ from matched cell types. For Capture C data^18,19^, we considered a loop overlapping if the ‘bait’ region listed the eGene and the ‘other’ region overlapped any significant eQTL SNP by bedtools intersect.

## Data and Code Availability

Summary results data files (eQTL all pairs files, etc) can be found at Zenodo DOI 10.5281/zenodo.18760368. FastQ files for raw sequencing reads and genotyping vcf are located in dbGAP, placement pending. Code used in this manuscript is at github bvoightlab/Tonsil_eQTL. All other data is available from the authors upon reasonable request.

## Acknowledgements

BFV is grateful for support by the NIH (DK138512, ES013508, and AI146026).

NR was supported by the NIH (AI179680, AI146026, AI184976) and the Jeffrey Modell Foundation Translational Research Award. Andrew Wells was supported by the NIH (AI154773, AI146026, DK122586).

## Author Contributions

BFV: Conceptualization, Investigation, Resources, Writing – Original Draft, Writing – Review and Editing, Visualization, Supervision, Project Administration. NR: Conceptualization, Investigation, Resources, Writing – Original Draft, Writing – Review and Editing, Visualization, Supervision, Project Administration. KL: Conceptualization, Methodology, Formal Analysis, Investigation, Data Curation, Writing – Original Draft, Writing – Review and Editing, Visualization. AW: Conceptualization, Investigation, Resources, Writing – Review and Editing, Visualization, Supervision, Project Administration. CLC: Conceptualization, Methodology, Investigation. SY: Methodology, Investigation, KZ: Resources, Methodology

