## Supplementary Figures for "Tonsillar expression quantitative trait loci verify and expand genetic contributors to childhood atopic diseases"

**Supplementary Table Descriptions**

Supplementary Table 1: eQTL Scoreboard. First tab is column header descriptions, other tabs correspond to tested cell types.

Supplementary Table 2: Age and Sex Interaction eQTL. Each tab is the output of tensorqtl for one cell type using the flags --interaction (for sex or age, as labelled) --best_only --mode cis_nominal

Supplementary Table 3: eQTL called on subsets of data based on age or sex. Each tab is the output of tensorqtl for one cell type using the flag --mode cis, parsed for pval_nominal < pval_nominal_threshold and qval < 0.05

Supplementary Table 4: GWAS Traits used for Colocalization. Contains summary information for each GWAS tested for colocalization with our eQTL.

Supplementary Table 5: Summary of Tonsil eQTL Colocalization Results. All tested colocalizations, with lead GWAS SNP, cell type, gene, PP0-PP4, and conditional PP4.

Supplementary Table 6: Approximate Bayes Factor Credible Sets.

Supplementary Table 7: Summary of DICE Colocalization Results. All tested colocalizations, with lead GWAS SNP, cell type, gene, PP0-PP4, and conditional PP4.

Supplementary Table 8: Details about eGenes identified as colocalizing with pediatric asthma.

Supplementary Table 9: DESEQ2 log2 fold changes for all tested genes.

Supplementary Table 10: DESEQ2 log2 fold changes for significant genes called across data subsets based on age or sex. First table describes columns, each other tab corresponds to one celltype-pair/subset combination (ie naive B vs GC B cells in male and female data subsets). Full results for these datasets available on Zenodo.

Supplementary Table 11: Regions based on GWAS leads used for colocalization.

**Files available on Zenodo**

allpairs/[celltype]_all.cis_qtl_pairs.txt.gz

files are tensorQTL output from the command "--mode cis_nominal", converted to plaintext from the original parquet file types

columns are

phenotype_id	variant_id	tss_distance	af	ma_samples	ma_count	pval_nominal	slope	slope_se

credsets/[celltype]_credsets.txt.gz

files are the results of approximate Bayes Factor credible set mapping

columns are

cell_type	gene_name	ENSG_ID chr	pos	a1	a2	pval_nominal	slope	slope_se	PIP

DE_subsets/[celltype1]_[celltype2]_DESEQ_results_[analysis]_lfc_het.txt

analyses are 4paired_vs_5paired and Mpaired_vs_Fpaired

correspond to supplementary figures 4 & 5

columns are

ENSG	gene_name	gene_biotype	chr	start	end	F_only_lfc	F_only_lfcSE	F_only_padj	M_only_lfc	M_only_lfcSE	M_only_padj	QQp	I2	interest	lfc_delta

eQTL_subsets/[celltype]_[analysis].cis_qtl.txt

files are tensorQTL output from the command "--mode cis"

analyses are male, female, lt4 (4 or younger) and gt5 (5 or older)

columns are

phenotype_id	num_var beta_shape1	beta_shape2	true_df pval_true_df	variant_id	tss_distance	ma_samples	ma_count	af	pval_nominal	slope	slope_se	pval_perm	pval_beta	qval	pval_nominal_threshold

gcts/[celltype]_readcount.gct.gz and gcts/[celltype]_tpm.gct.gz

collected output from RNAseq processing pipeline

in gct format

**Supplementary Figures**

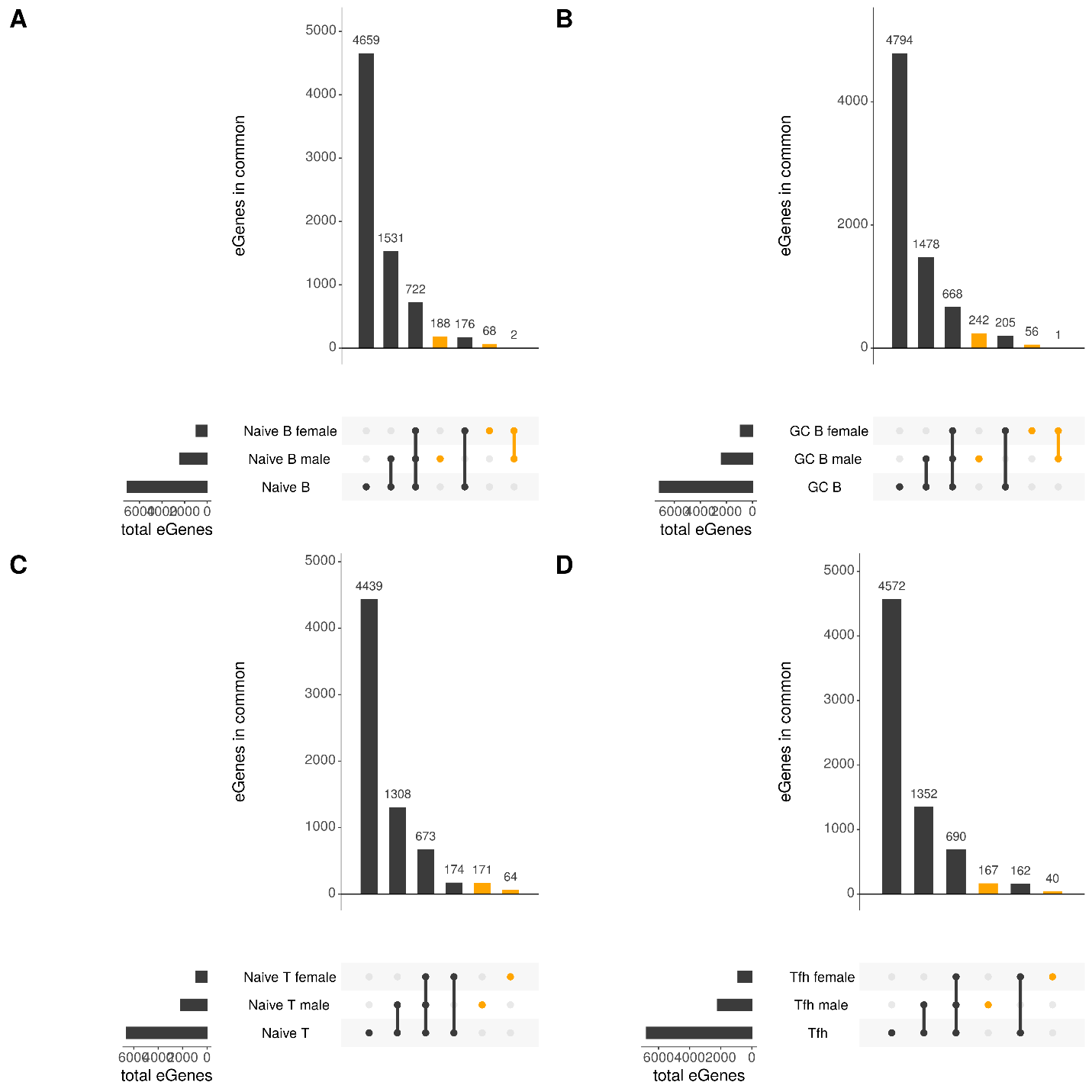

Supplementary Figure 1: UpSet plots comparing eGenes of sex based subsets of data.

UpSet plots showing the overlap between significant genes. eGenes present in a sex based subset only are highlighted in orange. Data shown separately for each cell type: A) naïve B, B) GC B, C) naïve T, and D) Tfh.

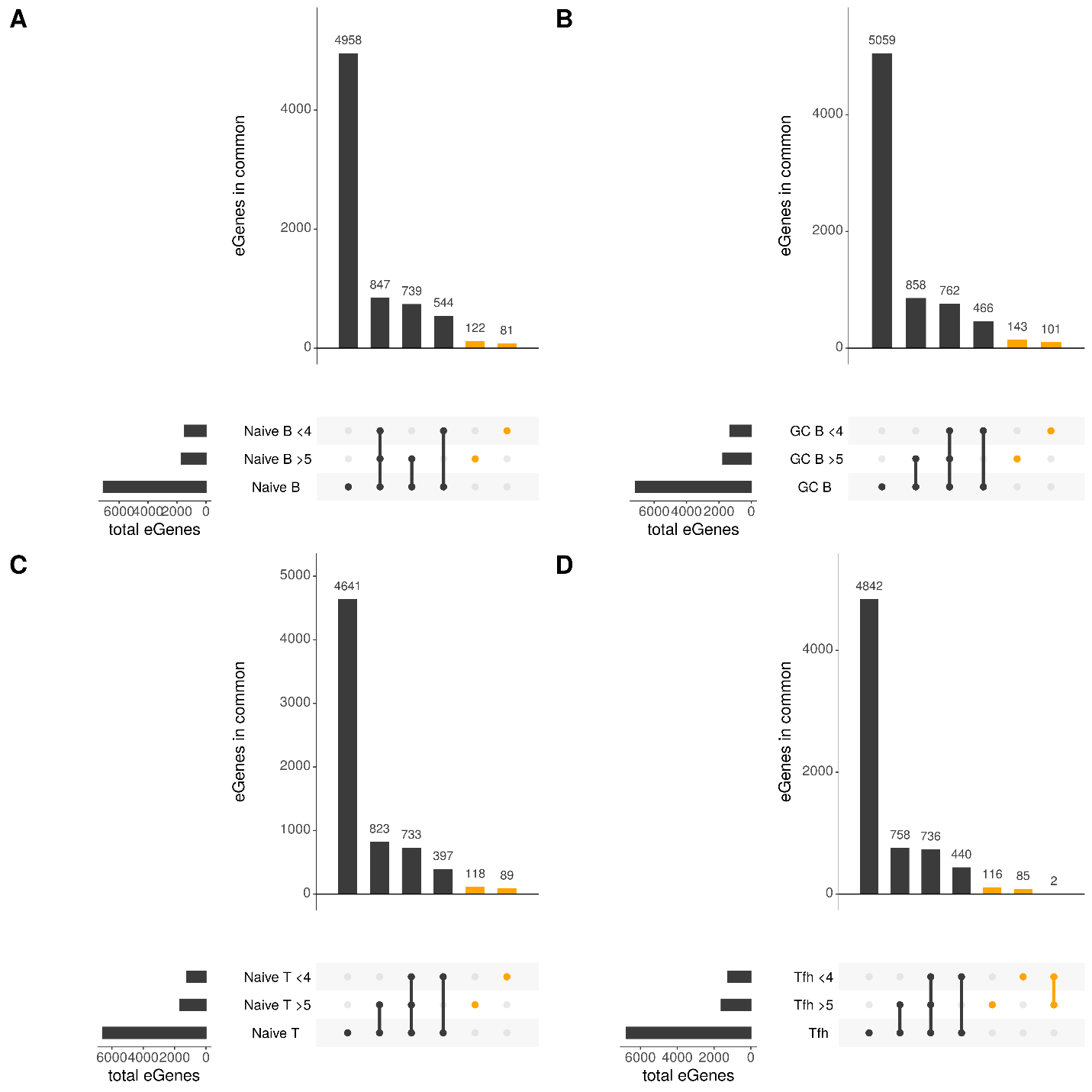

Supplementary Figure 2: UpSet plots comparing eGenes of age based subsets of data.

UpSet plots showing the overlap between significant genes. eGenes present in a age based subset only are highlighted in orange. Data shown separately for each cell type: A) naïve B, B) GC B, C) naïve T, and D) Tfh.

A

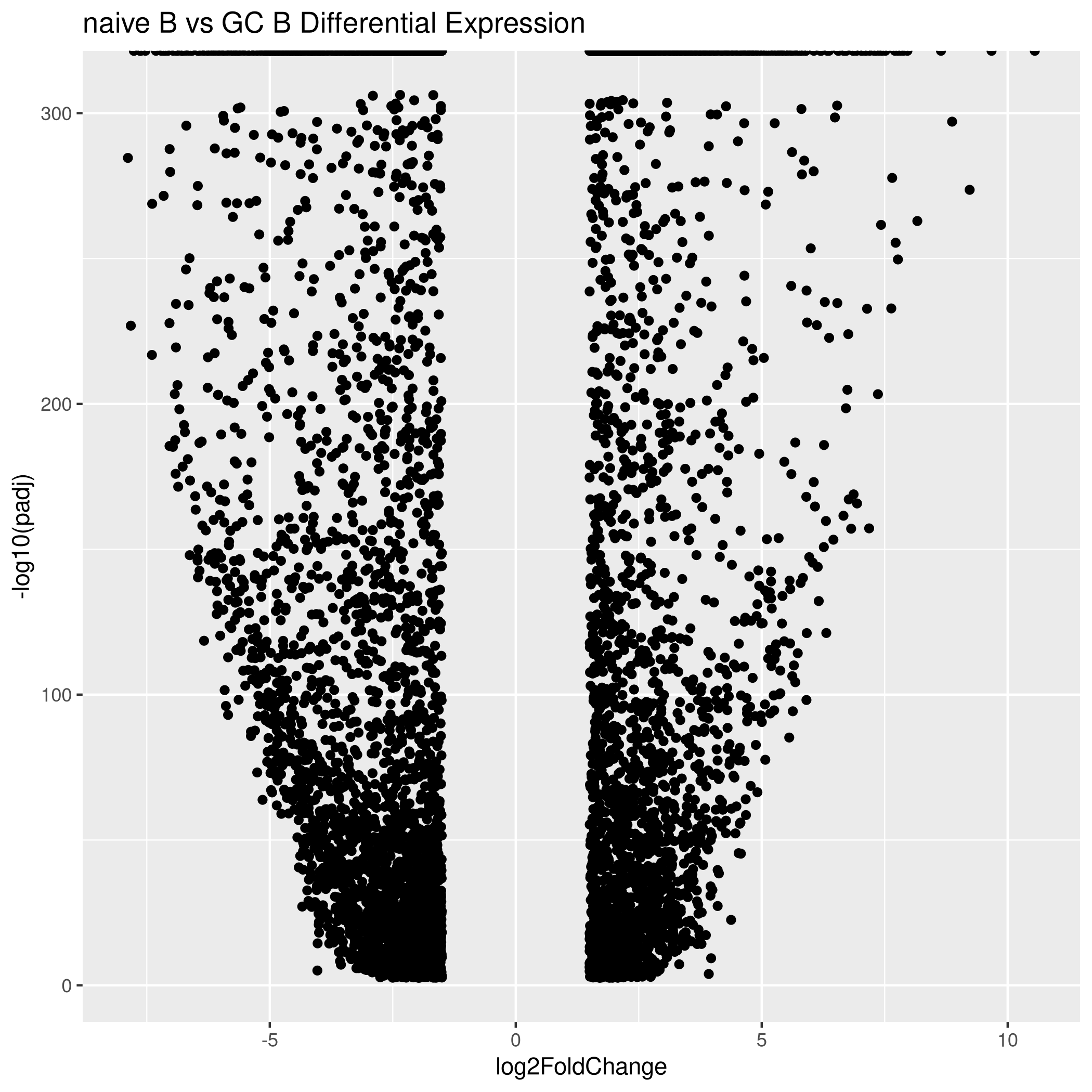

B

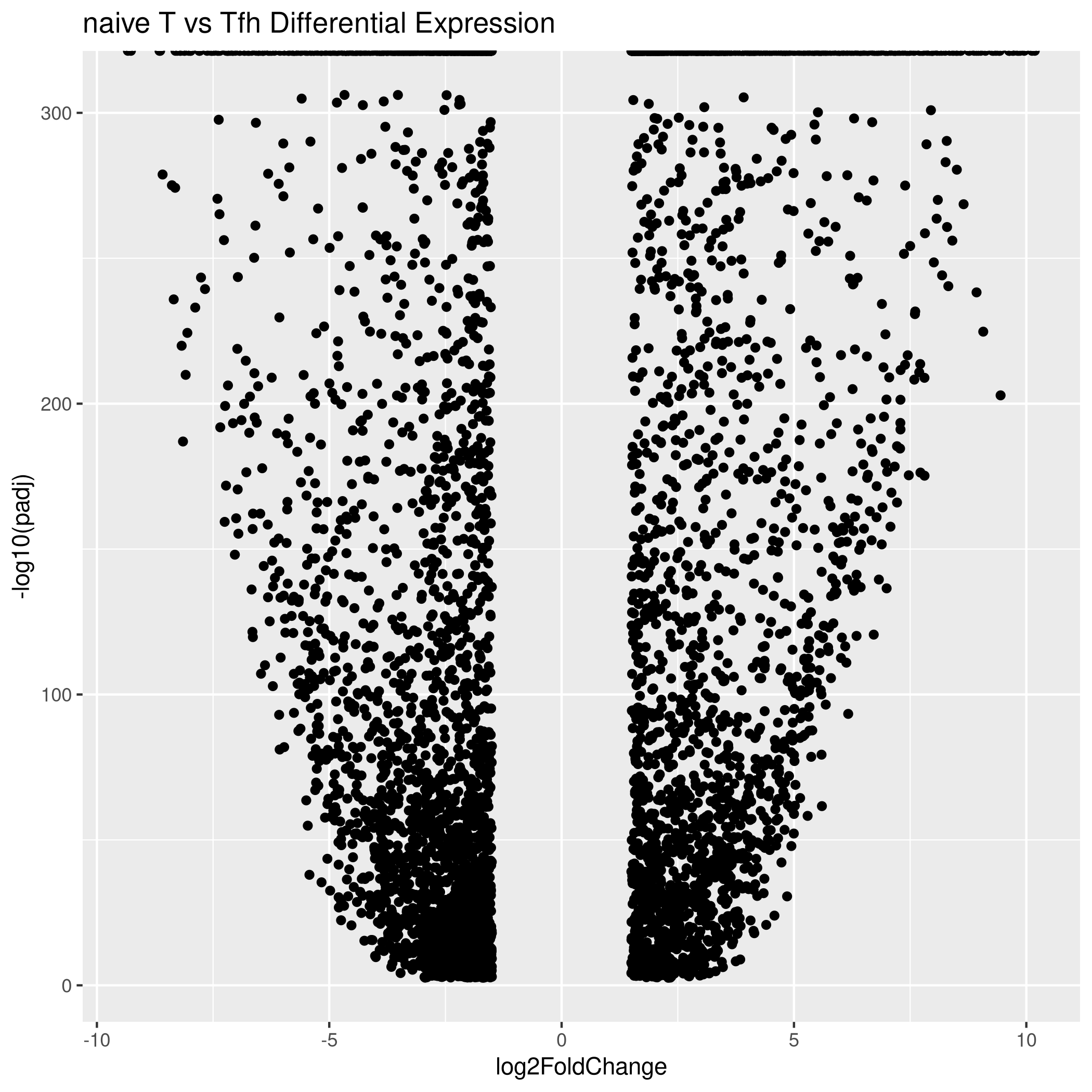

C

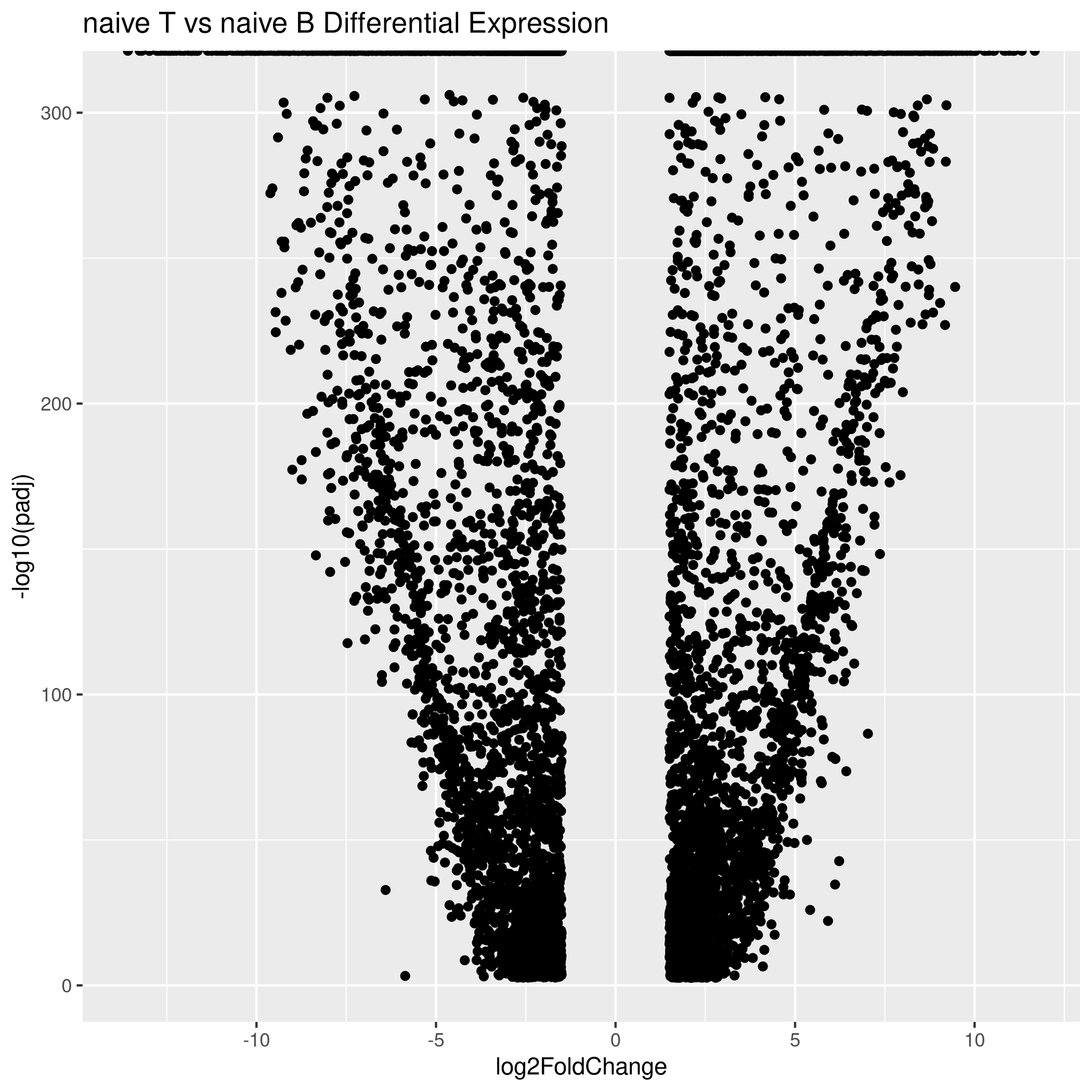

D
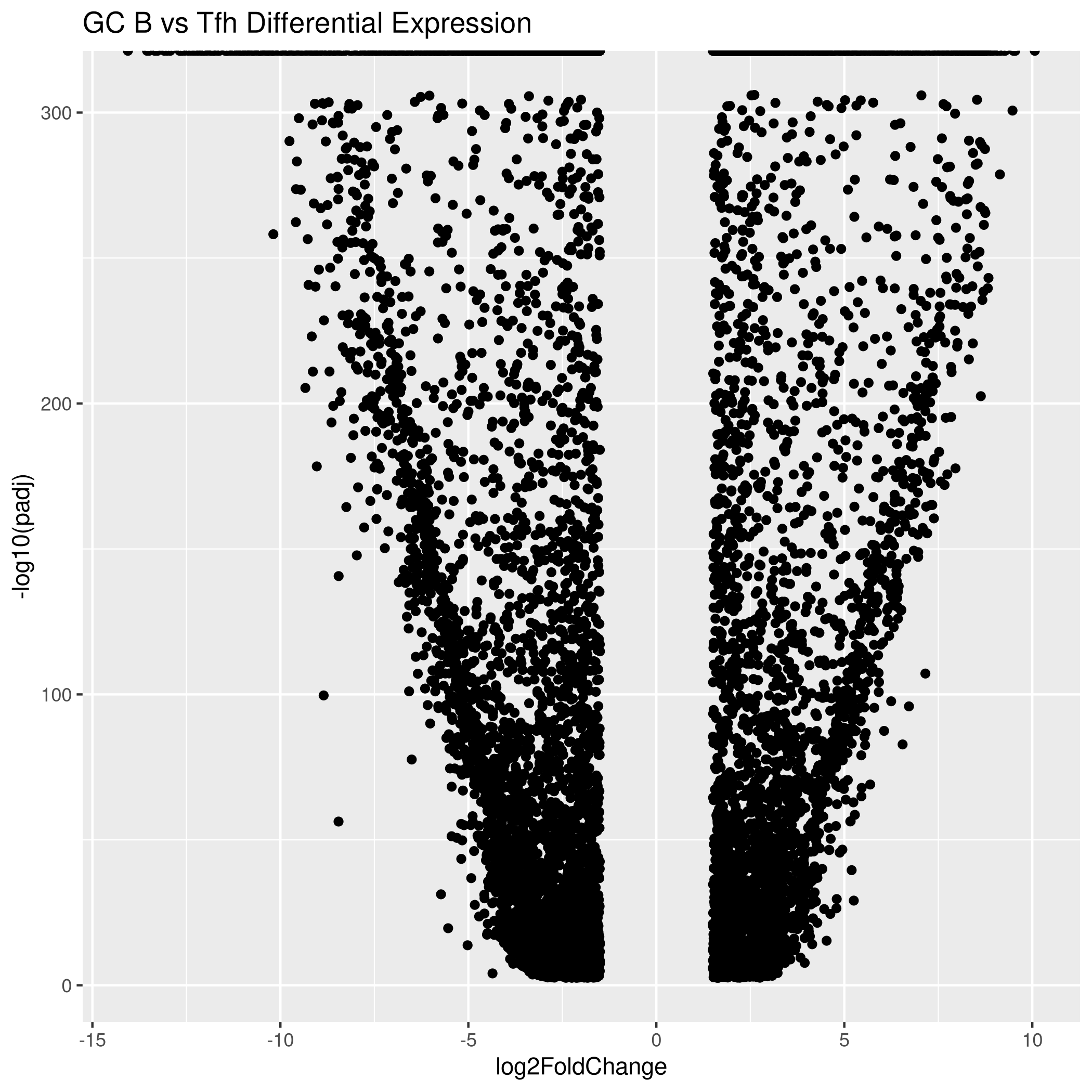

Supplementary Figure 3: Volcano plots showing differentially expressed genes.

Volcano plots with log2 fold change along the x-axis and -log10(adjusted p value) on y-axis. We only assessed relevant cell type combinations: A) B cell, naïve vs germinal center, B) T cell, naïve vs follicular helper, C) naïve B vs naïve T, and D) GC B vs Tfh.

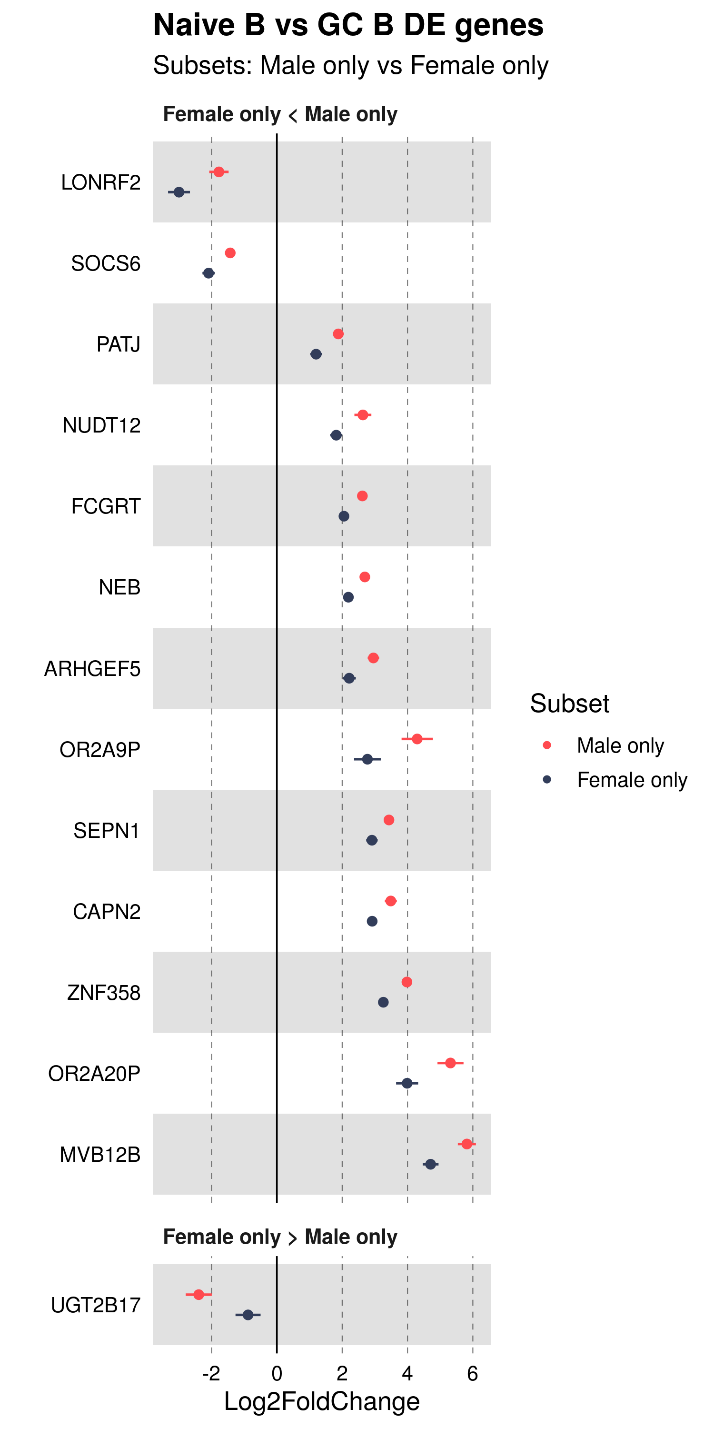

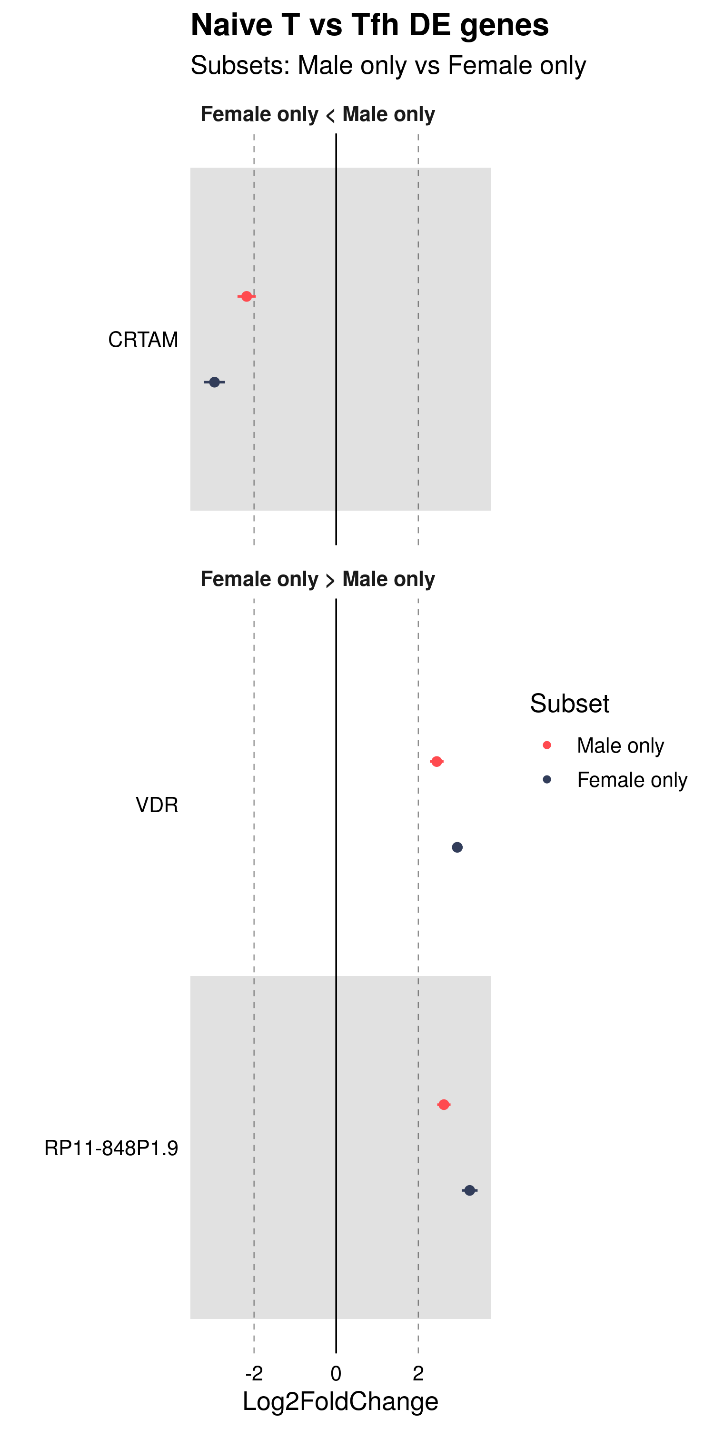

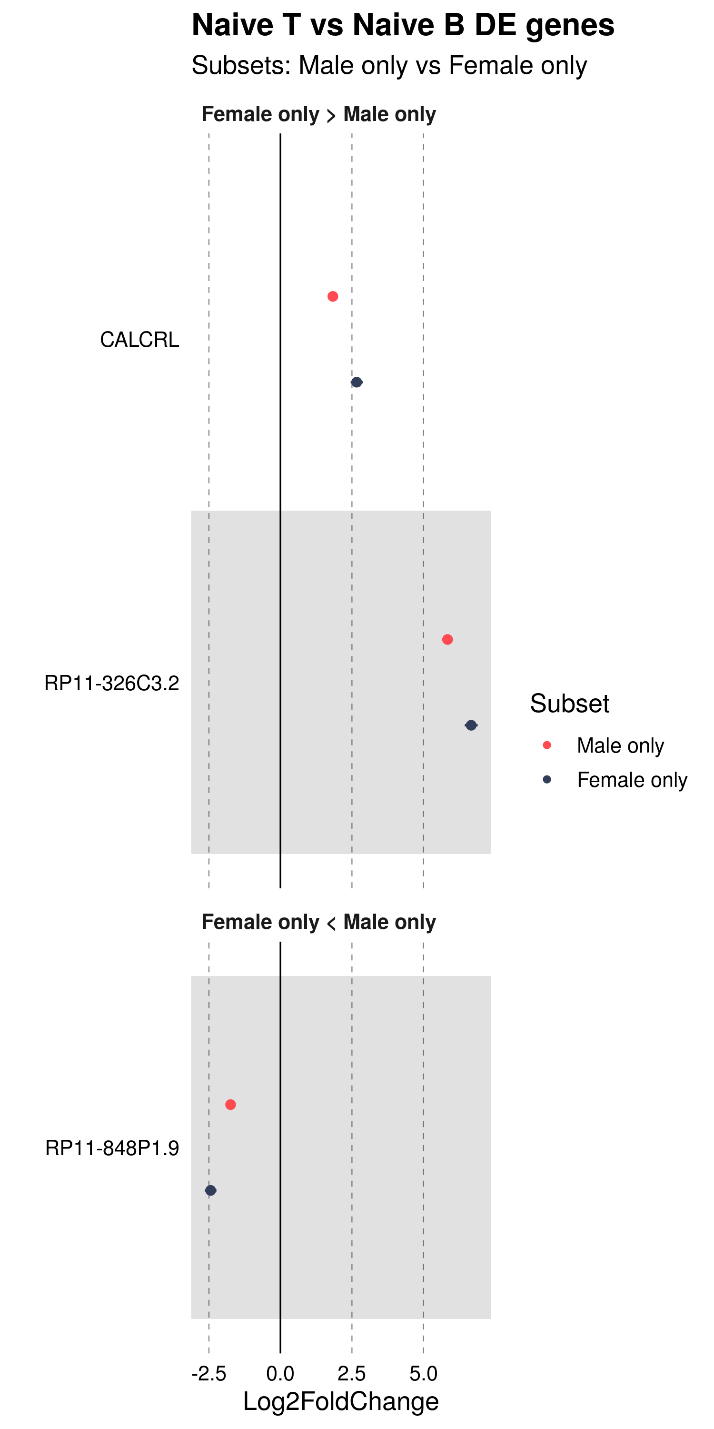

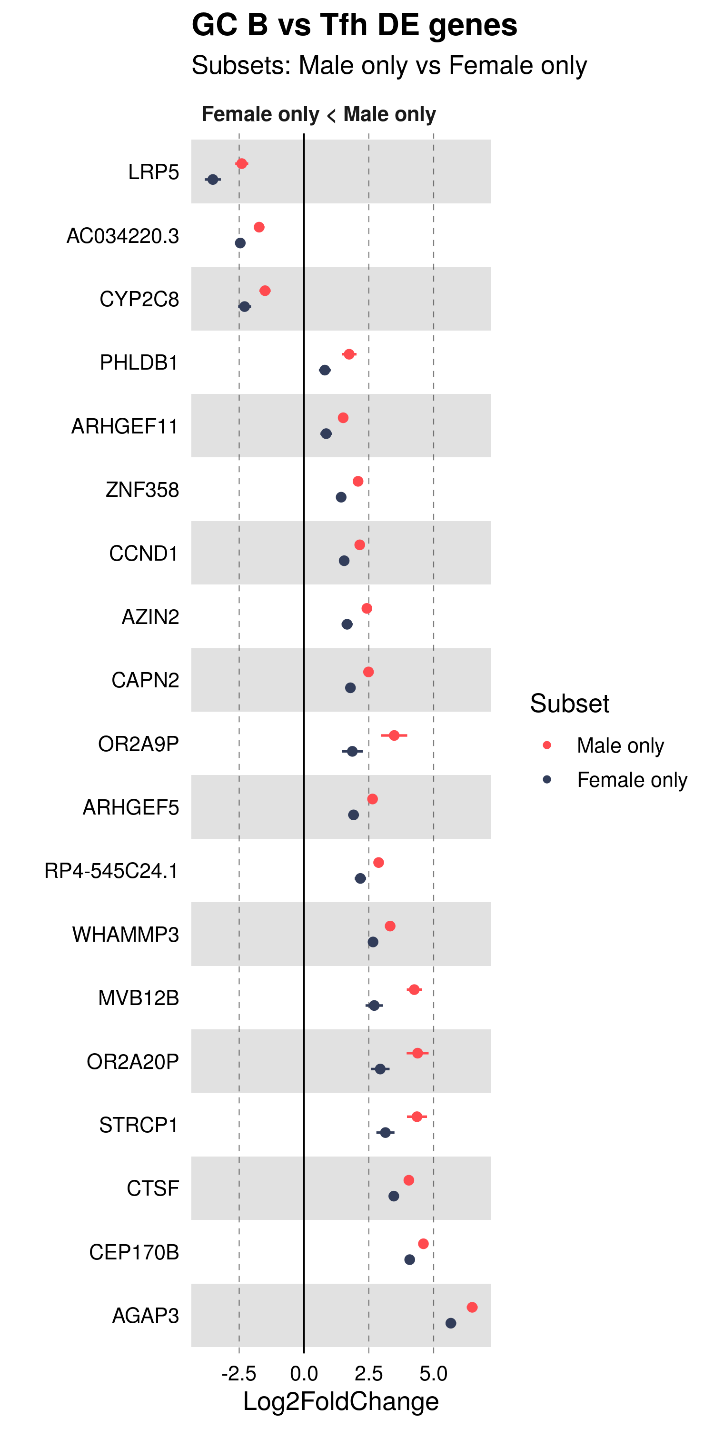

Supplementary Figure 4: Forest plots showing genes with heterogeneous log fold change between sex based subsets.

Forest plots with log_2_ fold change along the x-axis and genes along y-axis, with LFC of male subset in red and female subset in black. Top panel includes genes with lower LFC in male samples, bottom shows genes with lower LFC in female samples. We only assessed relevant cell type combinations: A) B cell, naïve vs germinal center, B) T cell, naïve vs follicular helper, C) naïve B vs T, and D) GC B vs Tfh.

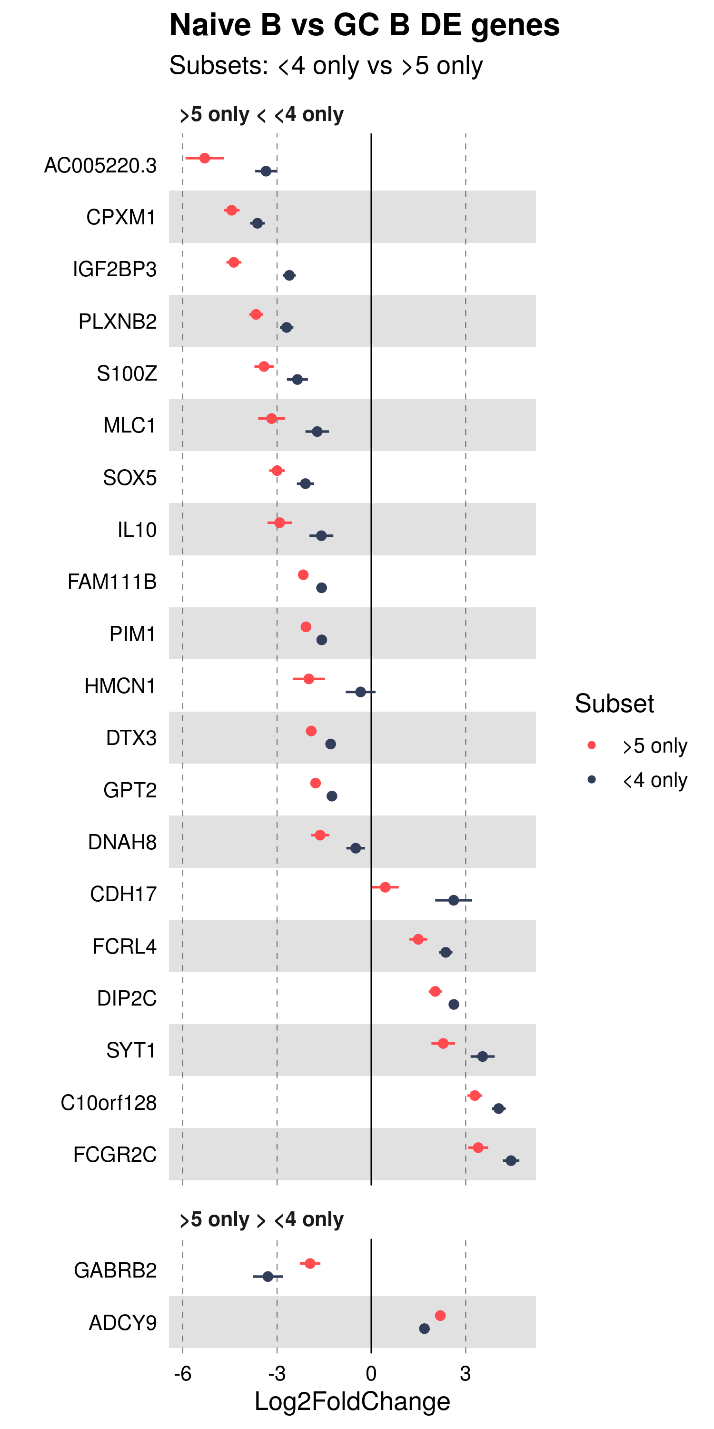

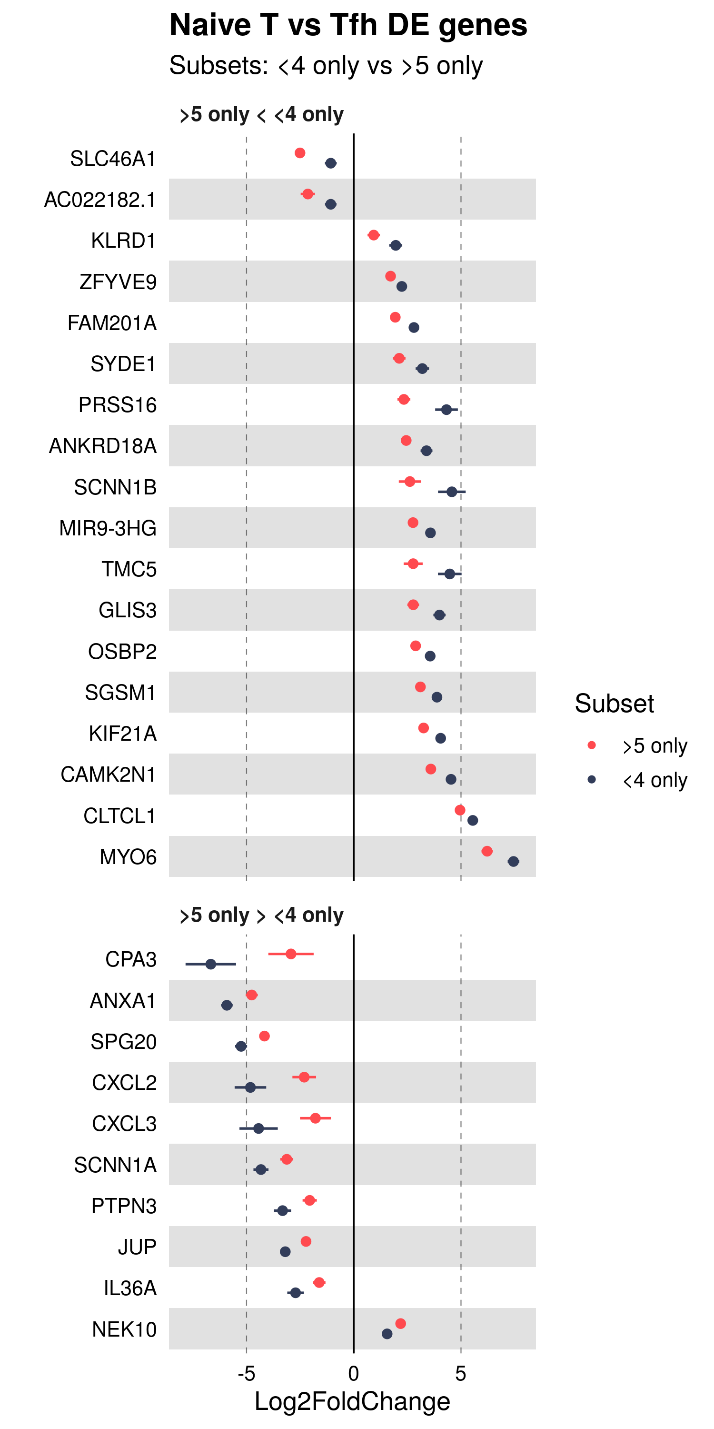

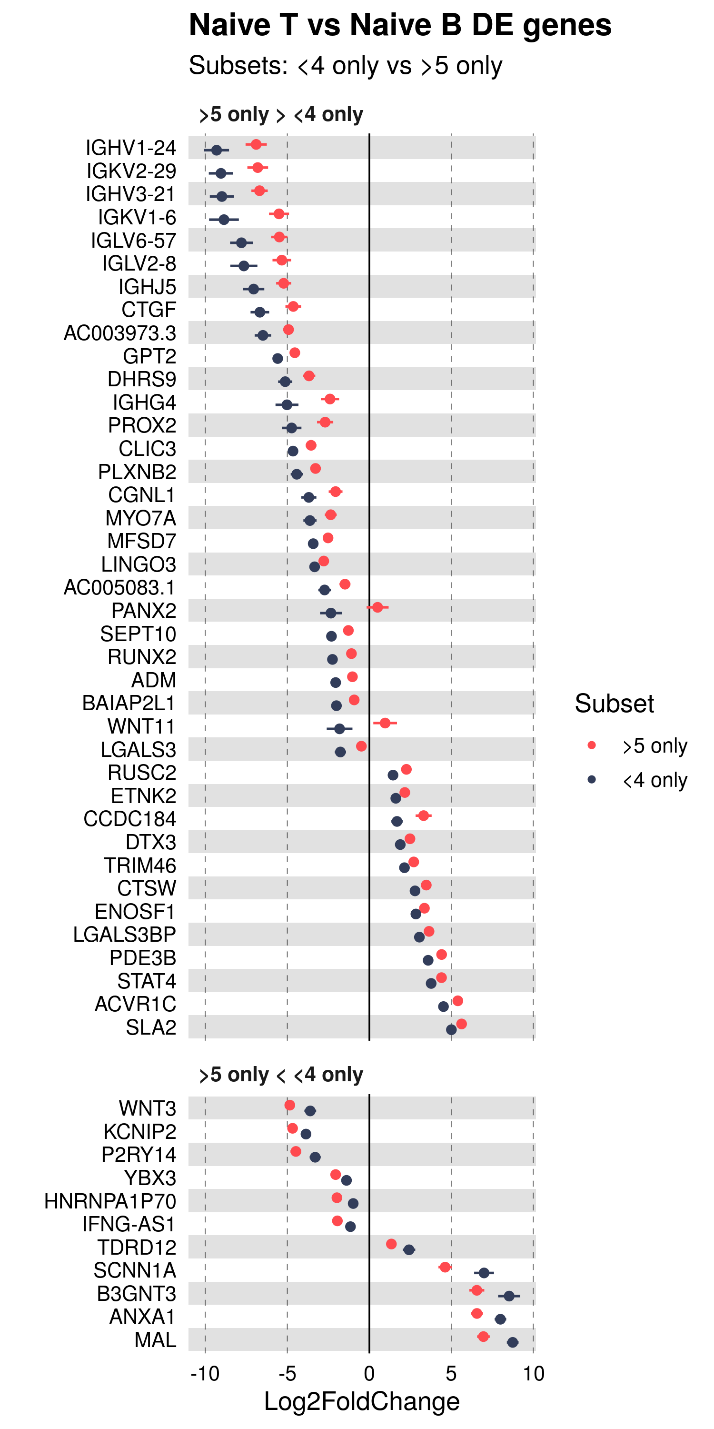

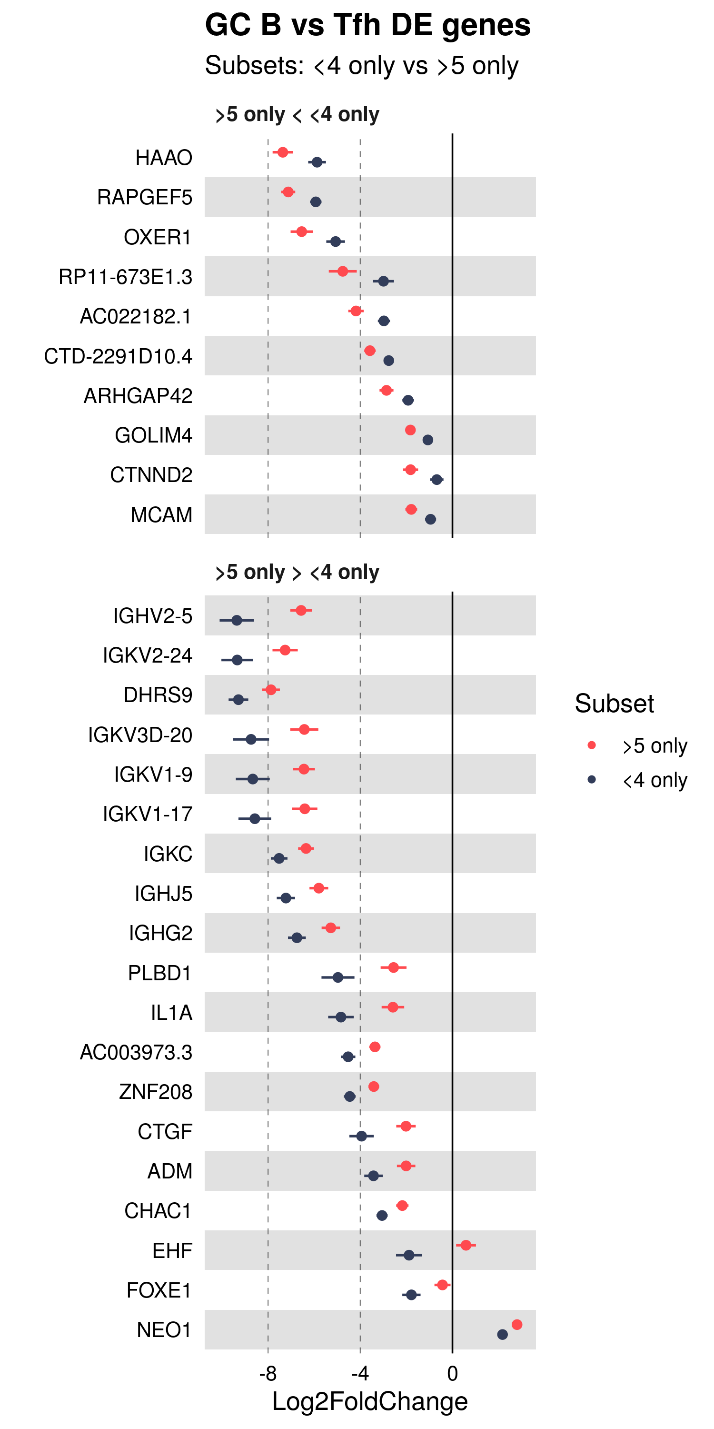

Supplementary Figure 5: Forest plots showing genes with heterogeneous log fold change between age based subsets.

Forest plots with log_2_ fold change along the x-axis and genes along y-axis, with LFC of older subset in red and younger subset in black. Top panel includes genes with lower LFC in younger samples, bottom shows genes with lower LFC in older samples. We only assessed relevant cell type combinations: A) B cell, naïve vs germinal center, B) T cell, naïve vs follicular helper, C) naïve B vs T, and D) GC B vs Tfh.

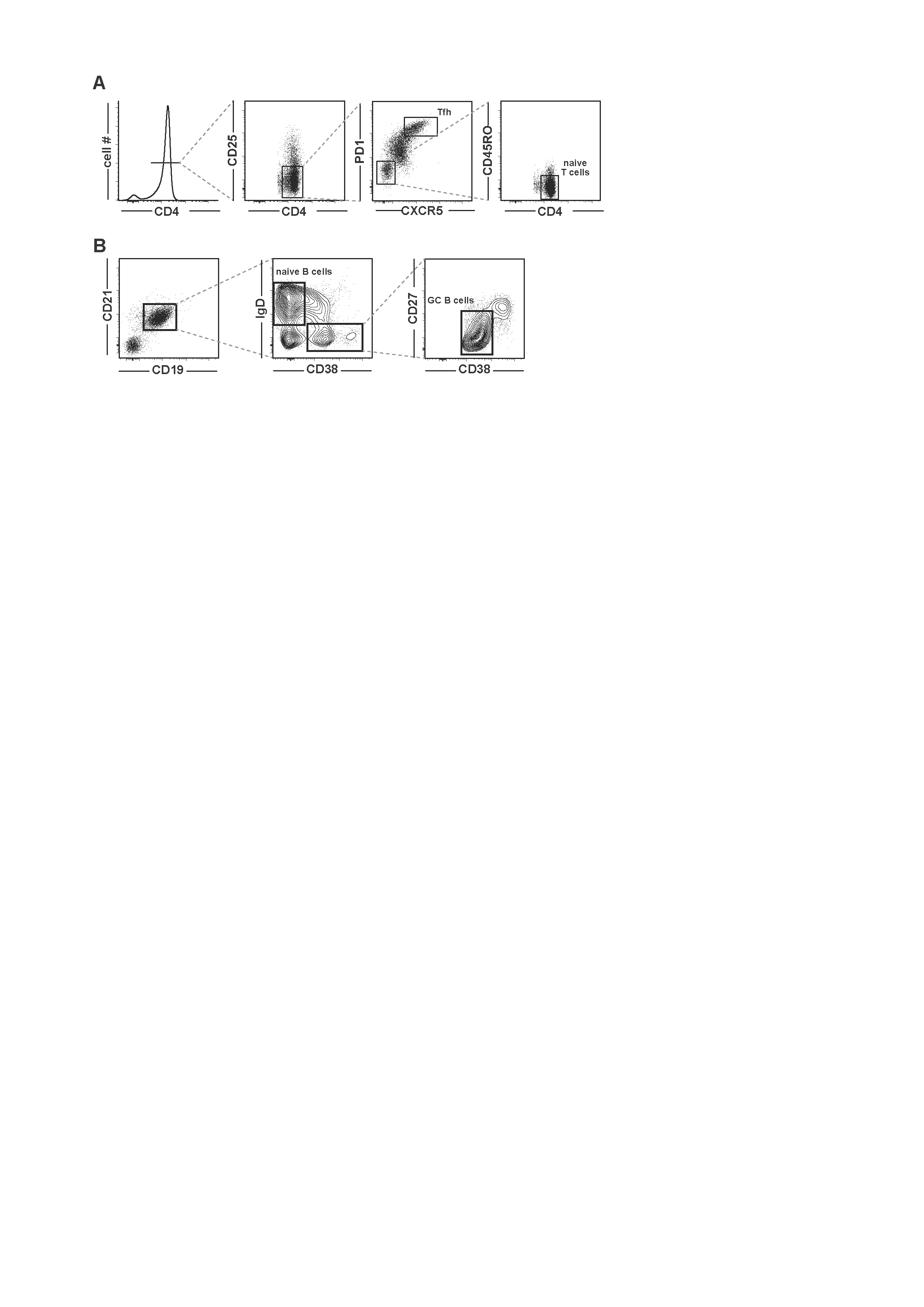

Supplementary Figure 6: Gating Strategy.

Gates used to sort A) Tfh and naïve T cells B) naïve B and GC B cells.

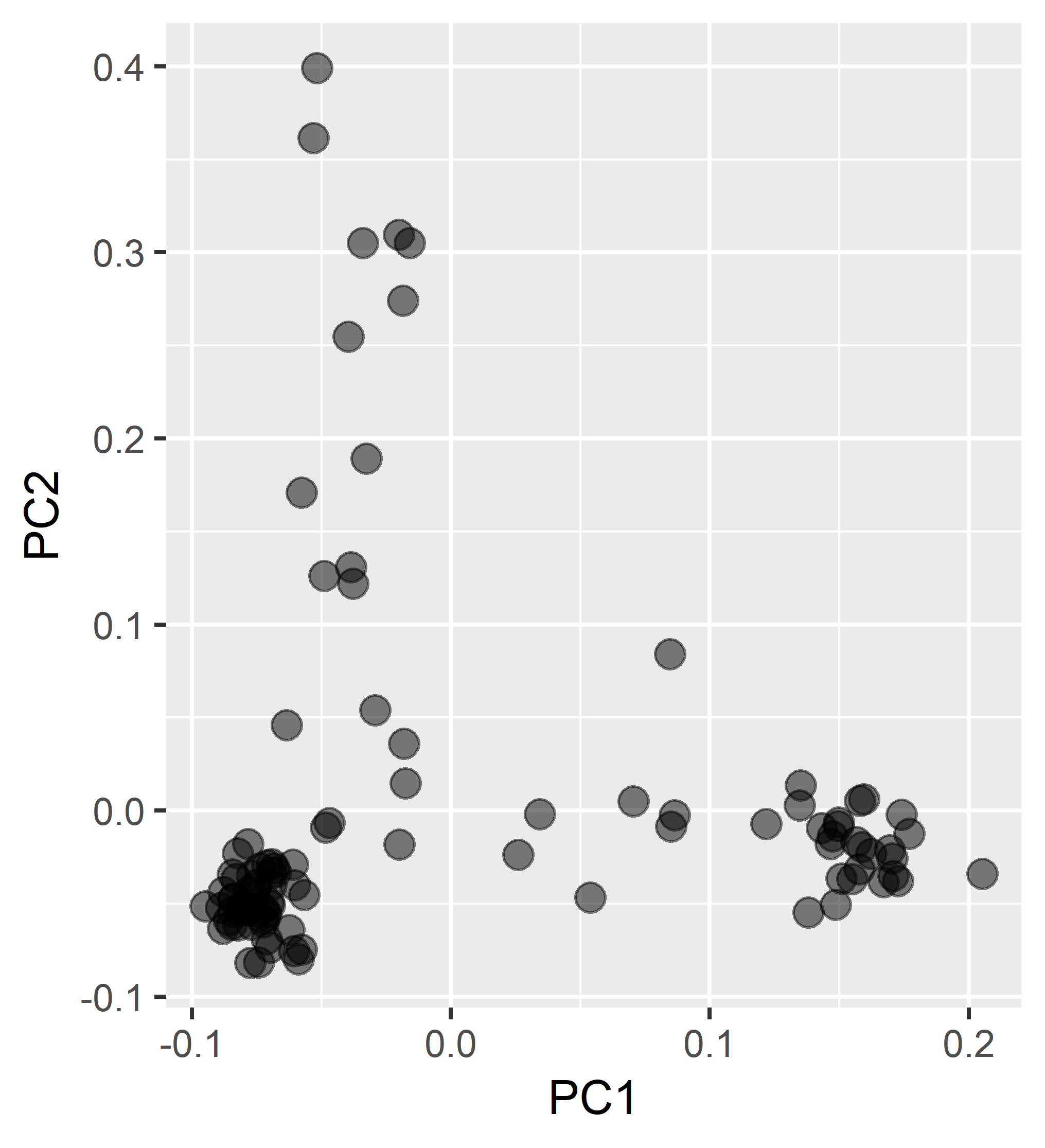

Supplementary Figure 7: Genotype Principal Components.

Plot shows PC1 and PC2, separating major ancestry components into the familiar triangle pattern, with EUR-like patients at the base and EAS-like and AFR-like patients along the two axes.
